# The effects of insecticide seed treatments on green peach aphid *Myzus persicae* (Sulzer) (Homoptera: Aphididae) parasitism by *Aphidius colemani* Viereck (Hymenoptera: Aphidiidae) and predation by *Mallada signatus* (Schneider) (Neuroptera: Chrysopidae)

**DOI:** 10.1101/2021.03.07.434302

**Authors:** Samantha Ward, Ary A. Hoffmann, Maarten Van Helden, Paul A. Umina

## Abstract

The green peach aphid, *Myzus persicae* (Sulzer) (Homoptera: Aphididae), is a major pest of *Brassica* L. species in Australia, where it can transmit >100 viruses. Globally, this species has evolved resistance to 74 insecticides from numerous chemical groups. Although Integrated Pest Management (IPM) strategies are being implemented, chemical treatment remains the predominant method used to control aphids. Insecticide seed treatments are viewed as a softer alternative to chemical sprays and are widely used in Australian canola fields. The effects of imidacloprid, thiamethoxam, and a mixture of thiamethoxam & lambda-cyhalothrin canola seed treatments were investigated on the parasitoid, *Aphidius colemani* Viereck (Hymenoptera: Aphidiidae) and the predator, the green lacewing *Mallada signatus* (Schneider) (Neuroptera: Chrysopidae); both important natural enemies of *M. persicae*. The number of mummies formed by *A. colemani* on the untreated plants was lower than those formed on the thiamethoxam & lambda-cyhalothrin and imidacloprid treated plants. The number of *A. colemani* reared from mummies on thiamethoxam & lambda-cyhalothrin plants was higher than those reared from thiamethoxam and untreated plants. Significant effects of insecticide seed treatments were only noted for mummies produced while the parent parasitoids were on the plants, not for those mummies produced after their removal. This suggests seed treatment effects were immediate but not long lasting. Based on cumulative parasitoid survival days for two generations, *A. colemani* exposed to thiamethoxam & lambda-cyhalothrin and imidacloprid treatments had a greater fitness than those exposed to the thiamethoxam and untreated controls, possibly due to the phenomenon of insecticide hormoligosis. Despite the treatment effects observed, we did not detect any behavioural differences in *M. persicae* or *A. colemani. Mallada signatus* were not negatively affected by feeding on *M. persicae* on insecticide seed treated plants, suggesting they are more tolerant of seed treatments than *A. colemani*. The findings from this study provide a useful platform for further experimentation on the effects of seed treatments on natural enemies of *M. persicae*.

## 1. Introduction

The generalist aphid, the green peach aphid, *Myzus persicae* (Sulzer) (Homoptera: Aphididae), is an important pest of *Brassica* L. species in temperate regions of the world (Cole, 1997). It is particularly damaging to field crops, feeding on over 400 species of plants in 40 different families and transmitting over 100 plant viruses (Blackman and Eastop, 2000, Kennedy et al., 1962). Since the early 1950s, insecticide use has been the main method for suppressing *M. persicae* populations (DeBach, 1974, Desneux et al., 2004, Gullan and Cranston, 1994, Hardin et al., 1995). This has contributed to *M. persicae* evolving resistance to more than 70 insecticides, from numerous chemical groups including carbamates, neonicotinoids, organophosphates, and pyrethroids (Bass et al., 2014, de Little et al., 2017, Umina et al., 2014). Although several agricultural systems utilize biological control as part of their Integrated Pest Management (IPM) strategies, chemical control is still predominantly relied upon to manage *M. persicae* in grain crops globally (Bass et al., 2014, Voudouris et al., 2017).

A potential issue associated with the use of insecticides is their toxicity to non-target beneficial arthropods, which can result in secondary pest outbreaks (El-Ghar and El-Sayed, 1992, Croft, 1990, Hardin et al., 1995, Roubos et al., 2014). It can be difficult to predict the side effects of insecticides, due to the many interacting variables (Jepson et al., 1990). This becomes more challenging when considering insecticides applied directly to crop seeds; with the efficacy of chemically treated seeds on multiple trophic levels often debated (Walters, 2013). Seed treatments involve placing an insecticide around the seed, with the chemical spreading through the plant systemically. Chemical sprays, however, can cover whole soil or plant surfaces, thus potentially endangering more target organisms (Dewar and Denholm, 2007).

Pyrethroids have long been used as foliar sprays to control aphids yet their overuse in the last decade has resulted in knockdown resistance (*kdr*; caused by an alteration in the target site of action), and so neonicotinoid seed treatments were used as an alternative (Dewar and Foster, 2017). Neonicotinoids began as seed treatments with the registration of imidacloprid in 1994 (Jeschke et al., 2011), quickly becoming the most widespread of all insecticides used as seed treatments (Douglas and Tooker, 2015, Huang et al., 2015, Miao et al., 2014). Although both causing paralysis and eventually death, neonicotinoids are nicotinic acetylcholine receptor (nAChR) allosteric modulators that act on the nerves, whereas pyrethroids are sodium channel modulators (Fernandes et al., 2016, IRAC, 2015). In addition to this, pyrethroids are known to possess repellent action towards arthropods (Burden, 1975, He et al., 2008).

Irrespective of exposure route, pyrethroids and neonicotinoids can produce a variety of effects on non-target insects. Imidacloprid has produced lethal effects on *Aphidius colemani* Viereck (D’Ávila et al., 2018). Along with another neonicotinoid thiamethoxam, imidacloprid has been recorded to produce lethal effects on *Aphidius gifuensis* Ashmead (Ohta and Takeda, 2015), predatory bugs and other parasitoids (Prabhaker et al., 2011). When exposed to thiamethoxam, detrimental effects have been recorded for multiple beneficial organisms, such as coccinellid larvae (Moscardini et al., 2015, Moser and Obrycki, 2009), multiple aphid predators (Seagraves and Lundgren, 2012), and green lacewings (*Chrysoperla carnea* (Stephens)) (Gontijo et al., 2014). Furthermore, this chemical can cause sublethal effects, reducing the proportion of female offspring produced by the parasitoid *Lysiphlebus testaceipes* (Cresson) (Moscardini et al., 2014). Yet, imidacloprid has been found to have negligible effects on predators when applied as a seed treatment (Epperlein and Schmidt, 2001) and no significant long-term detrimental effects on aphid natural enemies (Krauter et al., 2001). There have been a huge number of studies demonstrating the negative effects of pyrethroid chemicals on non-target species. Positive effects have also been detected with pyrethroids, for example the fecundity of the diamondback moth (*Plutella xylostella* (L.)) increases with exposure to fenvalerate (Fujiwara et al., 2002, Sota et al., 1998). Ideally, insecticides should be selective, highly toxic to pests but not to other organisms (Roubos et al., 2014).

In Australia, canola (*Brassica napus* L.) was first commercially cultivated in the late 1960s but was not grown widely around the country until the early 1990s (Colton and Potter, 1999). Similar to many countries, Australian canola is almost exclusively sown as insecticide-treated seed with a view to reducing pest threats during the early crop establishment period. *Myzus persicae* is one of the three major aphid pests infesting canola in Australia (Gu et al., 2007). Due to their ability to transmit turnip yellows virus (TuYV), *M. persicae* can cause yield losses of up to 50% (Berlandier, 2004). *Myzus persicae* often inhabit emerging canola seedlings which are particularly vulnerable at the cotyledon damage (Moens and Glen, 2002). Increasing insecticide resistance and environmental concerns highlight the importance of incorporating biological control into pest management strategies (Holloway et al., 2008, Umina et al., 2019).

Waterhouse and Sands (2001) list the natural enemies of *M. persicae* in Australia; ten species of predator, six parasitoids, eight hyperparasitoids, and six fungi. An important species is *A. colemani* (Hymenoptera: Braconidae; Aphidiidae), a pan-tropical species of parasitoid, widely distributed in Africa, Asia, Australia, South America, and southern Europe, that parasitizes Aphididae, including *M. persicae* (Starý, 1975). It is commercially available for the biological control of aphids (Grasswitz, 1998, Jones et al., 2003). Chrysopid species (green lacewings) are included among the most important aphidophagous predators (Pappas et al., 2011), and their use as a biological control agent for aphids has been documented for over 250 years (Senior and McEwen, 2001). Most chrysopid adults are not predaceous and feed on nectar, pollen and/or aphid honeydew (Pappas et al., 2011). *Mallada signatus* (Schneider) (Neuroptera: Chrysopidae) is native to Australia and New Zealand (Smithers, 1988) and produced commercially to control numerous crop pests (Simmons and Gurr, 2004).

The aims of this paper are a) to understand the direct (lethal) effects and indirect (sublethal) effects of insecticide seed treatments on *A. colemani* and *M. signatus* when exposed to *M. persicae* that have fed on insecticide-treated canola seedlings, and b) to draw comparisons between the impacts of canola seed treatments commercially relevant in Australian canola: 350g/l thiamethoxam (Cruiser^®^ 350FS), 210g/l thiamethoxam & 37.5g/l lambda-cyhalothrin (Cruiser^®^ Opti), and 600g/l imidacloprid (Gaucho^®^ 600).

## 2. Methodology

### 2.1. Seed treatments

Untreated ATR Stingray canola seeds were coated with one of three chemical treatments, using a Hege 11 seed treater (Wintersteiger, Ried im Innkreis, Austria) to produce the following:

1. 600g/l imidacloprid, at a rate of 400ml/100kg (commercially available in Australian canola, marketed as Gaucho^®^ 600)
2. 210g/l thiamethoxam & 37.5g/l lambda-cyhalothrin, at a rate of 1000ml/100kg (commercially available in Australian canola, marketed as Cruiser^®^ opti)
3. 350g/l thiamethoxam, at a rate of 600ml/100kg (marketed as Cruiser^®^ 350FS in Australian cereals)

The imidacloprid and thiamethoxam & lambda-cyhalothrin treatments are registered as seed treatments for Australian canola, however the thiamethoxam-only treatment is not, but was tested here to identify which, if any, active ingredient of thiamethoxam & lambda-cyhalothrin affects beneficial organisms. The rates of thiamethoxam were matched between treatments. 10-12 seeds of each treatment (in addition to untreated ‘control’ seeds) were planted within plastic pots (100 mm × 100 mm × 75 mm), in an unfertilised, non-sterilised, premium grade potting mix (with high soluble nitrogen content and water storing granules) (Table 1). These were placed within a controlled temperature (CT) room at 22°C (+/- 3°C), ∼40% relative humidity (R.H.) and a 16 light (L):8 dark (D) photoperiod. Each pot stood in a petri dish and was watered sparingly three times a week, for two weeks. Watering of pots was closely regulated to ensure overwatering did not occur and cause insecticide treatments to leach out of the soil. Each treatment group was allocated a different Bugdorm insect rearing cage (4F4590 series, 475 × 475 × 930 mm, Australian Entomological Supplies Pty Ltd., Bangalow, NSW, Australia) to avoid contamination of insects.

**Table 1:** Canola treatment groups for each trial and number of replicates undertaken, with Insecticide Resistance Action Committee (IRAC) chemical grouping (Sparks and Nauen, 2015).

| **Commercial name** | **Active ingredient/s** | **Class (IRAC chemical group)** | **Rate** | **Trial** | **Replicates** |
| --- | --- | --- | --- | --- | --- |
| *Gaucho® 600* | Imidacloprid | Neonicotinoid (4A) | 400ml/100kg | 1a | 6 |
|  |  |  |  | 1b | 8 |
|  |  |  |  | 2a | 7 |
|  |  |  |  | 2b | 10 |
| *Cruiser® Opti* | Thiamethoxam & lambda-cyhalothrin | Neonicotinoid (4A) & pyrethroid (3A) | 1000ml/100kg | 1a | 6 |
|  |  |  |  | 1b | 8 |
|  |  |  |  | 2a | 7 |
|  |  |  |  | 2b | 10 |
| *Cruiser® 350FS* | Thiamethoxam | Neonicotinoid (4A) | 600ml/100kg | 1a | 6 |
|  |  |  |  | 1b | 8 |
|  |  |  |  | 2a | 7 |
|  |  |  |  | 2b | 10 |
| *Untreated* | - | - | - | 1a | 6 |
|  |  |  |  | 1b | 8 |
|  |  |  |  | 2a | 7 |
|  |  |  |  | 2b | 10 |

### 2.2. Insects

A population of *M. persicae* was obtained from a tomato crop (*Solanum lycopersicum* L.) in Bendigo, Victoria, Australia (−36.5645, 144.81) on 3^rd^ November 2017. A colony was established in the laboratory on bok choy (*Brassica rapa subsp. chinensis*) after purging any signs of parasitism. Genotyping of this colony revealed a single multi-locus clone, which is known to possess resistant alleles for *MACE* and super-*kdr*, as well as amplification of the *E4* esterase gene and an increased copy number of the P450, CYP6CY3; these have been closely linked to phenotypic resistance to carbamates, pyrethroids, organophosphates, and neonicotinoids, respectively (de Little et al., 2017). This resistant clone was chosen due to the high mortality observed in an insecticide-susceptible clone when fed on insecticide treated canola seedlings during pilot studies.

*Myzus persicae* were maintained on untreated canola plants for several generations prior to this study. *Aphidius colemani* (provided by Biological Services, Loxton, SA, Australia) were reared on susceptible *M. persicae* on canola plants. *Mallada signatus* (Bugs for Bugs, Toowoomba, QLD, Australia) were fed susceptible *M. persicae* on canola plants. For both beneficial insect species, diet was supplemented with a 20% honey solution and water, changed weekly. Additionally, the *M. signatus* colony was provided with bee pollen from untreated wildflowers (SaxonBee Enterprises, Gidgegannup, WA, Australia). All species were maintained within bug dorms, within a CT room maintained at 22°C (+/- 3°C), ∼60% R.H. and a 16L:8D photoperiod.

### 2.3. Experimental set up

Two weeks after sowing, the healthiest eight plants were selected (at the two-leaf stage), and the remaining plants removed from each pot. Each pot, still standing within a petri dish, was allocated to its own microcosm plastic container (102 mm × 108 mm × 200 mm), with mesh sides, and a lid with a mesh window, to allow for air flow. 100 *M. persicae* were added to each container. All plants continued to be watered sparingly three times a week, using a squeeze bottle directly into the pot, so as not to disturb any aphids. After 96 hours, the aphids were counted for a 0-DAT recording.

### 2.4. *Trial 1a –* Aphidius colemani

At 0-DAT, six mated female *A. colemani* were released into each microcosm container, after incubation at 4°C for five minutes (Table 1). Mating was assumed, as parasitoid sexes were stored together prior to the experiment, with mating usually occurring almost immediately after emergence (Starý, 1970). This experiment was conducted in a CT room at 22°C (+/- 3°C). After 24 hours, all living parasitoids were collected and stored separately in petri dishes, lined with filter paper and containing a wick dipped in a 20% honey solution (Starý, 1970). Duration of seed treatment exposure was selected for parasitoids based on previous similar experiments undertaken, such as that by Carter (2013). Wicks were replaced every three days, or more often if dried out or showing signs of mould. Fifty *M. persicae* from the original colony (and not exposed to seed treatments) were then added to each petri dish, which were maintained at 22°C (+/- 3°C), ∼60% R.H. and a 16L:8D. Survival of parasitoids was recorded daily to understand direct effects. These petri dishes containing the parental *A. colemani* were checked for mummies 11 days later. Mummies were identified by their engorged, golden/brown appearance, dissimilar to unparasitized aphids (Askew, 1971).

In addition, experimental plants were checked for mummies 10 days after the removal of parental *A. colemani*. Mummies were removed from leaves with a paintbrush (as in Buitenhuis et al. (2005)) and placed within individual petri dishes for rearing. All emerging first generation (F1) *A. colemani* (from petri dishes and from experimental plants) were removed and stored at -80°C. Petri dishes were checked daily, and mortality recorded. All dead parent *A. colemani* were removed and stored at - 80°C. A schematic of the experimental design is shown in Fig. 1.

**Figure 1:**
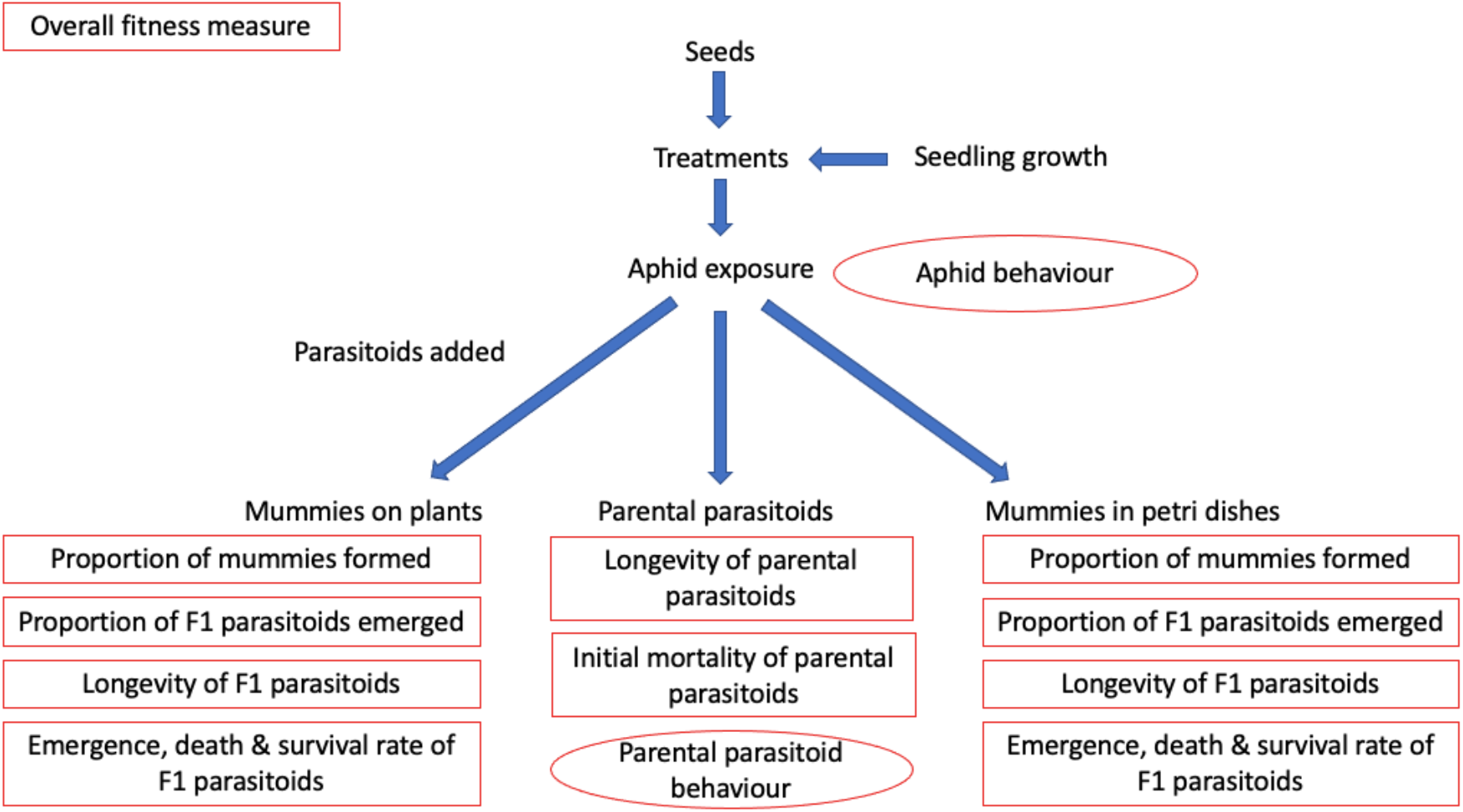
Schematic of *A. colemani* experiment, with data collected for trial 1a results boxed below each respective experimental pathway, and data collected for trial 1b results circled.

### 2.5. *Trial 1b - Aphid and* Aphidius colemani *behavioural experiment*

*Aphidius colemani* behaviour was investigated because significant treatment effects were detected within trial 1a. Behavioural experiments were undertaken to identify whether there were any treatment effects for either trophic level (pest and/or parasitoid), which could explain the significant results. This experiment was conducted in a CT room at 22°C (+/- 3°C). As per section 2.3., *M. persicae* were maintained on canola plants for 96 hours, after which time aphid numbers were counted to provide a 0-DAT count. The following day, twelve aphids were removed from each treatment and placed within individual petri dishes under a Leica MS5 microscope mounted with a Leica IC80 HD camera. Each aphid was video recorded for five minutes and their behaviour assessed using traits adapted from Bilodeau et al. (2013) (see Table S1).

For the parasitoid behavioural experiment, six mated female *A. colemani* were released onto each plant, as in trial 1a. Across all repeats, twelve parasitoids were removed from each treatment and placed individually within fresh petri dishes after 24 hours exposure. Along with each parasitoid, a naïve aphid (one that has not been in contact with parasitoids) from the same chemical treatment was added (Table 2). Aphids from different treatments (imidacloprid, thiamethoxam, and thiamethoxam & lambda-cyhalothrin) were also added to 12 unexposed *A. colemani* (Table 2). This was undertaken to determine that, if there were treatment effects on the interactions between parasitoid and pest, whether this was due to the aphids’ behaviour, rather than the parasitoids. *Aphidius colemani* were video recorded for five minutes and their behaviour assessed using traits adapted from Bilodeau et al. (2013) (see Table S1). A schematic of the experimental design is shown in Fig. 1, alongside trial 1a.

**Table 2:** Combinations of treatments for behavioural experiments.

| Aphid treatment | Parasitoid treatment | Replicates |
| --- | --- | --- |
| Untreated | - | 12 |
| Imidacloprid | - | 12 |
| Thiamethoxam | - | 12 |
| Thiamethoxam & lambda-cyhalothrin | - | 12 |
| Untreated | Untreated | 12 |
| Imidacloprid | Imidacloprid | 12 |
| Thiamethoxam | Thiamethoxam | 12 |
| Thiamethoxam & lambda-cyhalothrin | Thiamethoxam & lambda-cyhalothrin | 12 |
| Untreated | Imidacloprid | 12 |
| Untreated | Thiamethoxam | 12 |
| Untreated | Thiamethoxam & lambda-cyhalothrin | 12 |

### 2.6. *Trial 2a –* Mallada signatus

At 0-DAT, two 3^rd^ stage *M. signatus* larvae and two 1^st^ stage *M. signatus* larvae were released into each microcosm container (Table 1). After 24 hours, all living lacewings were collected and stored separately within petri dishes, lined with filter paper and containing a wick dipped in a 20% honey solution. This duration was chosen to match the exposure time of *A. colemani* within trial 1a and, as for the other trials, this experiment was conducted in a CT room at 22°C (+/- 3°C). One hundred *M. persicae* from the original colony (and not exposed to seed treatments) were added to each petri dish. These were replenished whenever numbers depleted. Wicks were replaced every three days, or more often if dried out or showing signs of mould. Once in adult form, bee pollen from untreated wildflowers (SaxonBee Enterprises, Gidgegannup, WA, Australia) was added to each petri dish, in place of *M. persicae* as a food source. All lacewings were maintained at 22°C (+/- 3°C), ∼60% R.H. and a 16L:8D. Petri dishes were checked daily for pupation, emergence and mortality. Dead *M. signatus* were removed and stored at -80°C within individual vials. Adult longevity was capped at 120 days, after which adult lacewings were removed and stored at -80°C. A schematic of the experimental design is shown in Fig. 2.

**Figure 2:**
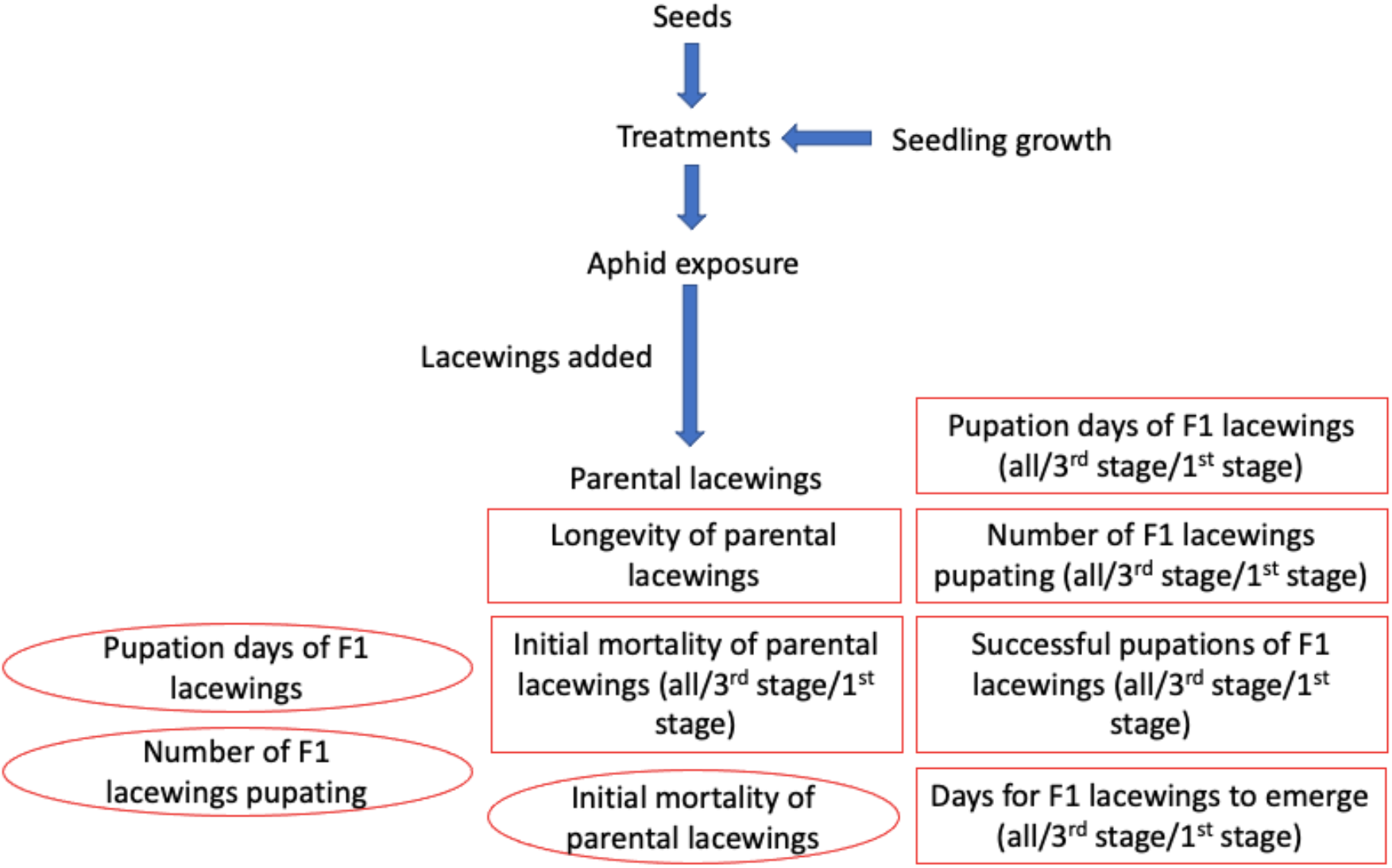
Schematic of *M. signatus* experiment, with data collected for trial 2a results boxed, and data collected for trial 2b results circled.

### 2.7. *Trial 2b –* Mallada signatus *pupation*

Due to high mortality in the 1^st^ stage *M. signatus* larvae in trial 2a, in part due to cannibalism among individuals (which is not uncommon (Duelli 1981)), a second experiment was conducted using one lacewing per container. This experiment was conducted in a CT room at 22°C (+/- 3°C). At 0-DAT, a single 1^st^ stage *M. signatus* larva was placed into a microcosm container with 100 *M. persicae* on untreated and seed treated plants (see Table 1). After 96 hours, all living lacewings were collected and stored separately within petri dishes, lined with filter paper and containing a wick dipped in a 20% honey solution. Lacewings were kept on the plants for 96 hours, as opposed to 24 hours in trial 2a, because no significant effects were detected during trial 2a. One hundred *M. persicae* from the original colony (and not exposed to seed treatments) were added to each petri dish as a food source and were replenished whenever numbers depleted. Wicks were replaced every three days, or more often if dried out or showing signs of mould. All lacewings were maintained at 22°C (+/- 3°C), ∼60% R.H. and a 16L:8D. Petri dishes were checked daily for pupation and once pupation occurred, pupae were removed and stored at -80°C within individual vials. A schematic of the experimental design is shown in Fig. 2, alongside trial 2a.

### 2.8. Mass spectrometry

Mass spectrometry was undertaken on canola plant material and *M. persicae* specimens from each treatment. This was undertaken by the Biotechnology & Synthetic Biology Group Land & Water at the Commonwealth Scientific and Industrial Research Organisation (Black Mountain, ACT, Australia) to determine the presence and concentration of imidacloprid, thiamethoxam, and clothianidin (a metabolite of thiamethoxam (Bredeson et al., 2015, Nauen et al., 2003)) (Table 3). Insect and plant samples were shipped to the facility on dry ice and freeze dried on arrival.

**Table 3:** A summary of samples tested for mass spectrometry analysis.

| Treatment | Trophic level | Number of specimens | Weight/s (mg) | Replicate/s |
| --- | --- | --- | --- | --- |
| Untreated | Pest | 200 aphids | 4.4-5.6 | 3 |
| Imidacloprid | Pest | 200 aphids | 5.2-5.6 | 3 |
| Thiamethoxam | Pest | 200 aphids | 3.8-4.4 | 3 |
| Thiamethoxam & lambda-cyhalothrin | Pest | 200 aphids | 4.6-5.8 | 3 |
| Untreated | Plant | 2 cotyledons | 2.7 | 1 |
| Imidacloprid | Plant | 2 cotyledons | 3.3 | 1 |
| Thiamethoxam | Plant | 2 cotyledons | 3.4 | 1 |
| Thiamethoxam & lambda-cyhalothrin | Plant | 2 cotyledons | 3.5 | 1 |

A Restek LC multiresidue pesticide standard #5 (Restek, 31976) was used to make standard curves for imidacloprid, thiamethoxam and its metabolite clothianidin. This was initially expressed in parts per billion (ppb), but due to high concentrations in the plant samples, the results were later expressed in parts per million (ppm). The standards were diluted in acetonitrile and covered four orders of magnitude in the range of 0.1 ppb to 1000 ppb (covering 0.1, 0.5, 1.0, 5.0, 10, 50, 100, 500 and 1000 ppb) and were run at the beginning and end of the analysis. The standard curve was linear up to 500 ppb (Fig. S1).

Samples were ground with a 6 mm stainless steel ball in 1 ml of 80% acetonitrile using a Qiagen TissueLyser at 30 Hz for three minutes, followed by incubation at 4°C for 30 minutes. Samples were centrifuged to pellet cell debris and the supernatant applied to an Agilent Captiva EMR Lipid media plate to clean the sample from major contaminants. The Captiva EMR Lipid media was previously tested to ensure the compounds of interest were not bound to the Captiva EMR media. The supernatant, 800 µl, was applied to the Captiva EMR media and the recovery of 600 µl which was dried down and resuspended in 100 µl prior to analysis.

Lambda-cyhalothrin was purchased from Sigma-Aldrich (St. Louis, MO, USA) and included in the mass spectrometry analysis, however it was unable to be detected at the levels within the plant matter or within the *M. persicae* in our study. Issues are often associated with the detectability of lambda-cyhalothrin (Gonçalves and Alpendurada, 2005).

### 2.9. Data analysis

All analyses were conducted using Minitab version 19.1.0.0 (Minitab, 2019). One-way ANOVAs were undertaken to analyse aphid population growth, for the aphid and parasitoid behavioural experiments, and to analyse rates of emergence, death and survival of parasitoids. Additional one-way ANOVAs were performed on log-transformed data to investigate the number of mummies formed per parasitoid. These were followed with Tukey pairwise post-hoc comparisons. GLMs were applied to analyse parent parasitoid survival over time. Finally, a cox regression analysis was undertaken to determine differences between the treatments for parent *A. colemani* survival over time.

To investigate the treatment effects on emergence/death rates of F1 parasitoids produced on untreated and seed treated plants, the data was transformed into cumulative days (i.e. ((‘number of parasitoids emerged on day 1’*1)+(‘number of parasitoids emerged on day 2’*2) and so on, with this total then divided by the total number of emerged parasitoids, and the computed emergence rates analysed with a one-way ANOVA. Death rates were computed in the same way as emergence rates except counting the cumulative number of parasitoids dying per day, divided by the total number of parasitoid deaths. To analyse F1 *A. colemani* longevity (survival rate) of those produced on the seed-treated or untreated plants, survival data was calculated as the ‘death rate’ subtracted from the ‘emergence rate’; treatment effects for this measurement were analysed using one-way ANOVAs.

In addition to the above, a total fitness measure for parasitoids was computed as another way to explore effects of the seed treatments. This consisted of cumulative parasitoid survival days, calculated by the following measure: (‘number of emerged parasitoids on day 1’*1) + (‘number of emerged parasitoids on day 2’*2)… + (‘number of emerged parasitoids on day 11’*11). This measure incorporated counts of all parent *A. colemani*, F1 *A. colemani* produced on the untreated or seed treated plants, and F1 *A. colemani* produced within the petri dishes. This is important from a biocontrol perspective because it provides a measure of the number of parasitoids available to parasitise eggs across the entire trial.

For lacewings, one-way ANOVAs were undertaken to analyse adult longevity, emergence time for lacewings and pupal duration in trial 2a. For trial 2b, pupal duration was also measured with a one-way ANOVA. These were followed by Tukey pairwise post-hoc comparisons. Treatment effects on initial mortality of lacewings and the number of successful lacewing pupations within trials 2a and 2b were analysed using contingency tests. The initial mortality and number and success of lacewings pupating were then compared for the different nymphal stages (1^st^ larval stage versus 3^rd^ larval stage lacewings) with paired t-tests. These tests were used to determine if the mean difference between the two sets of observations (1^st^ larval stage versus 3^rd^ larval stage lacewings) was zero.

## 3. Results

### 3.1. Mass spectrometry

Results for the mass spectrometry are presented in Table 4. The standard curves for each chemical tested are shown in Fig. S1. The slopes, intercept area and R^2^ values for each chemical are as follows: imidacloprid (slope = 0.97, intercept = 7.5, R^2^ = 0.998), thiamethoxam (slope = 0.92, intercept = 8.6, R^2^ = 0.997), and clothianidin (slope = 0.93, intercept = 7.3, R^2^ = 0.998).

**Table 4.**
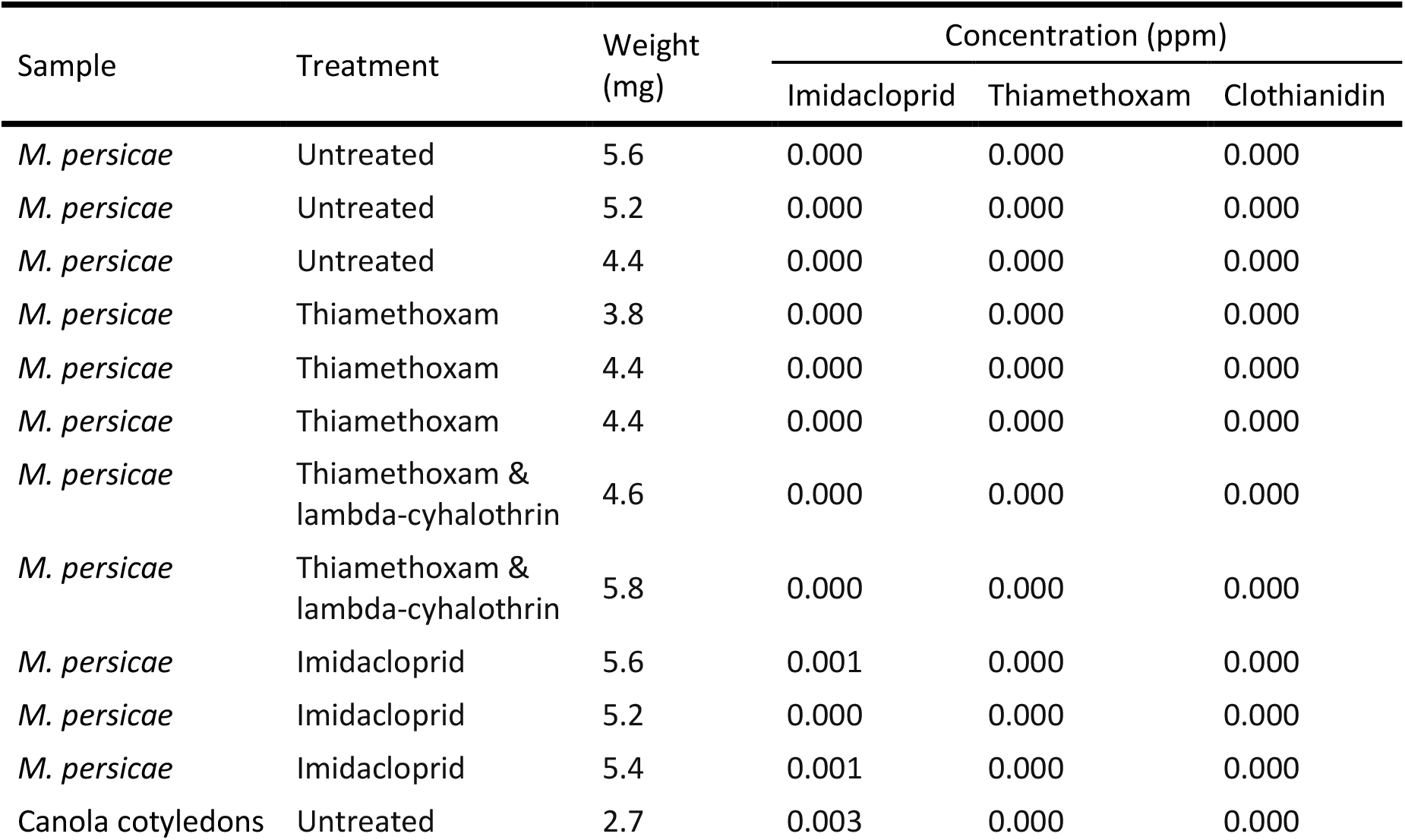
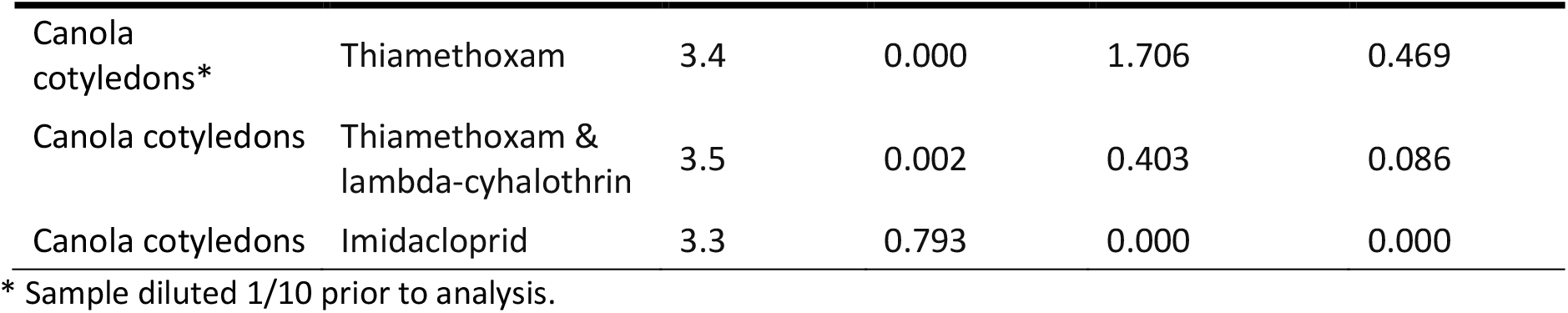
Final concentrations of imidacloprid, thiamethoxam, and clothianidin in plant matter and *M. persicae* from the various chemical treatments.

Imidacloprid and thiamethoxam were detected at high levels in their respective treated plants, with lower levels of thiamethoxam being detected in the thiamethoxam & lambda-cyhalothrin treatment when compared with the thiamethoxam-only treatment. As expected, clothianidin, the metabolite of thiamethoxam, was also present in the thiamethoxam and thiamethoxam & lambda-cyhalothrin treated canola seedlings. Imidacloprid was found within the untreated and thiamethoxam & lambda-cyhalothrin plant material, albeit in extremely very low quantities (Table 4).

Very low levels of imidacloprid were found within *M. persicae* taken from imidacloprid-treated canola seedlings. However, no thiamethoxam or clothianidin were found within aphids taken from thiamethoxam and thiamethoxam & lambda-cyhalothrin treated canola seedlings (Table 4).

### 3.2. Myzus persicae

Significant differences in the population growth of *M. persicae* were recorded after 96 hours exposure to seed treatments during the parasitoid experiment (trial 1a) (ANOVA, F_(3,56)_=19.72, p<0.001), and during the lacewing pupation experiment (trial 2b)) (ANOVA, F_(3,24)_=31.57, p<0.001), but not during the lacewing longevity experiment (trial 2a) (ANOVA, F_(3,36)_=2.56, p=0.078). The seed treated plants caused greater rates of *M. persicae* mortality for each succeeding trial. *Myzus persicae* populations grew during trial 1a at an average population increase of 64% for the untreated treatment, 72% for the imidacloprid treatment, 116% for the thiamethoxam treatment, and 160% for the thiamethoxam & lambda-cyhalothrin treatment (Fig. 3). During trial 2a, *M. persicae* populations had an average population increase of 56% for the untreated treatment, 30% for the imidacloprid treatment, 42% for the thiamethoxam treatment, and 76% for the thiamethoxam & lambda-cyhalothrin treatment (Fig. 3). Finally, in trial 2b, mean *M. persicae* population growth had a 32% increase for the untreated treatment, and the seed treatments showed population declines of 45% for the imidacloprid treatment, 30% for the thiamethoxam treatment, and 28% for the thiamethoxam & lambda-cyhalothrin treatment (Fig. 3).

**Figure 3:**
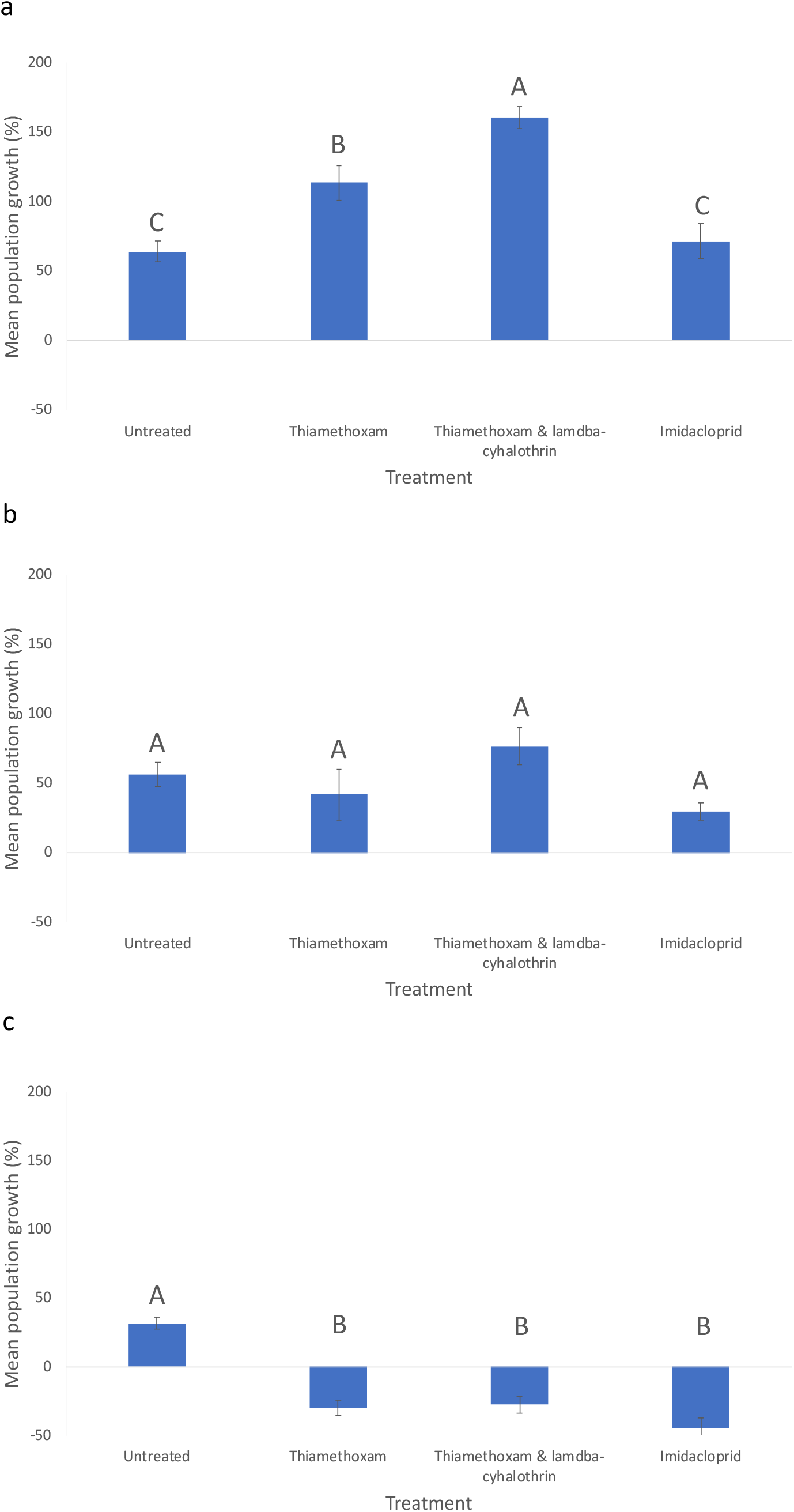
*M. persicae* population growth at 96 hours after introduction onto plants in a) trial 1, b) trial 2a, four months later, and c) trial 2b, another five weeks later. Letters represent significant differences between treatments within each trial. Error bars represent standard errors of the mean.

### 3.3. Aphidius colemani

Due to the differences between aphid numbers, shown in figure 3, mummies were calculated as a percentage of total number of aphids per treatment to standardise the analyses.

#### 3.3.1. Trial 1a – Survival of parental parasitoids

There was no difference detected between the seed treatments or untreated controls for parental *A. colemani* survival (number of parasitoids alive) after 24 hours (GLM: Treatment, F_(3,20)_=0.33; p=0.801; Fig. 4). A Cox regression analysis indicated no significant difference between the treatments for parental *A. colemani* survival (number of parasitoids alive) throughout time (days passed) (F_3, 19_=1.91, p=0.163, R^2^=23.13%; Fig. 4).

**Figure 4:**
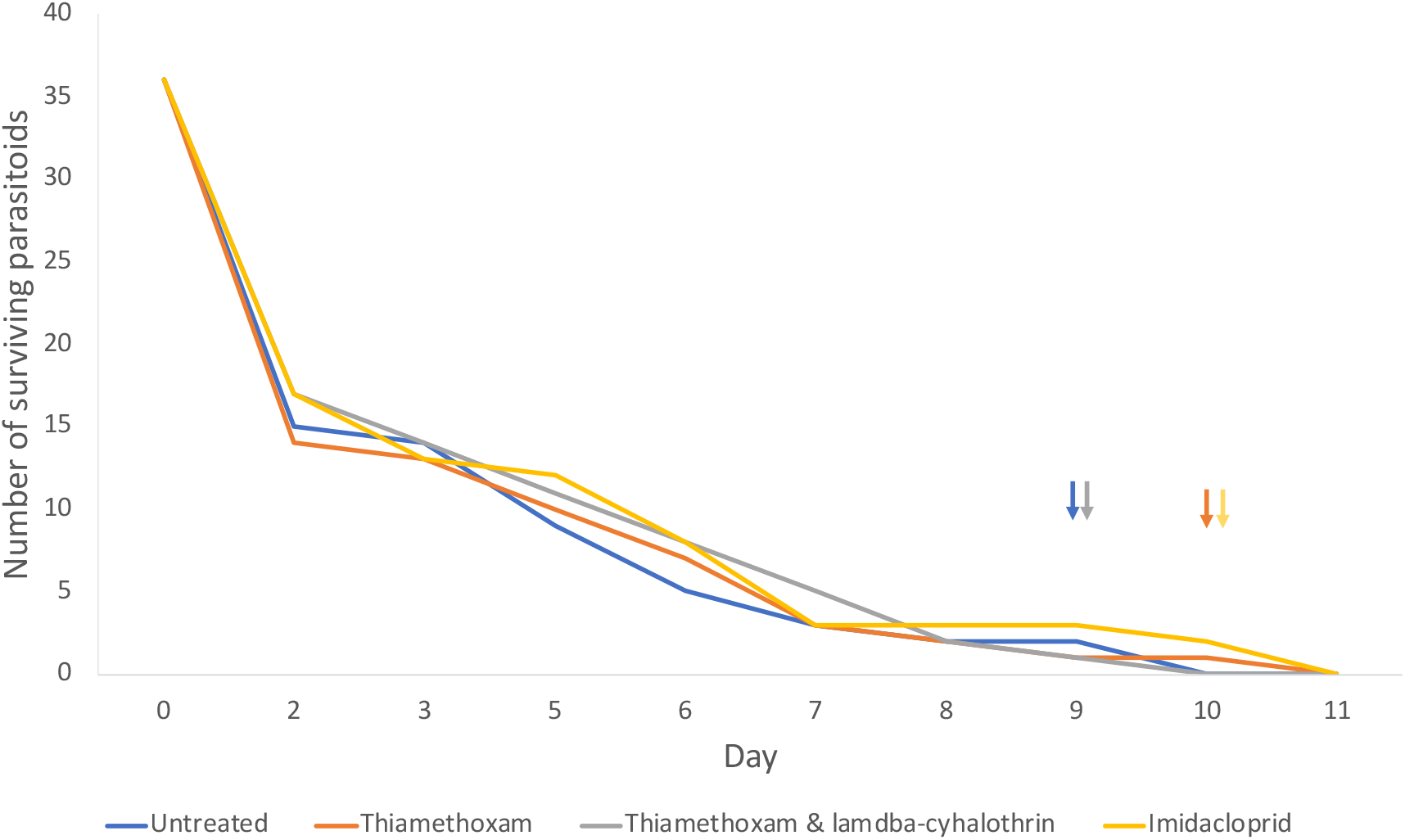
*A. colemani* survival for different seed treatments (arrows indicate last day of parasitoid survival for the colour coded treatment).

#### 3.3.2. Trial 1a – Mummies collected from plants

Fewer aphid mummies were produced in the untreated control canola, resulting in an overall treatment effect (F_3,19=_7.66, p<0.001; Fig. 5). Post-hoc tests indicate the number of mummies formed on untreated plants were significantly lower than those formed on thiamethoxam & lambda-cyhalothrin and imidacloprid treated plants.

**Figure 5:**
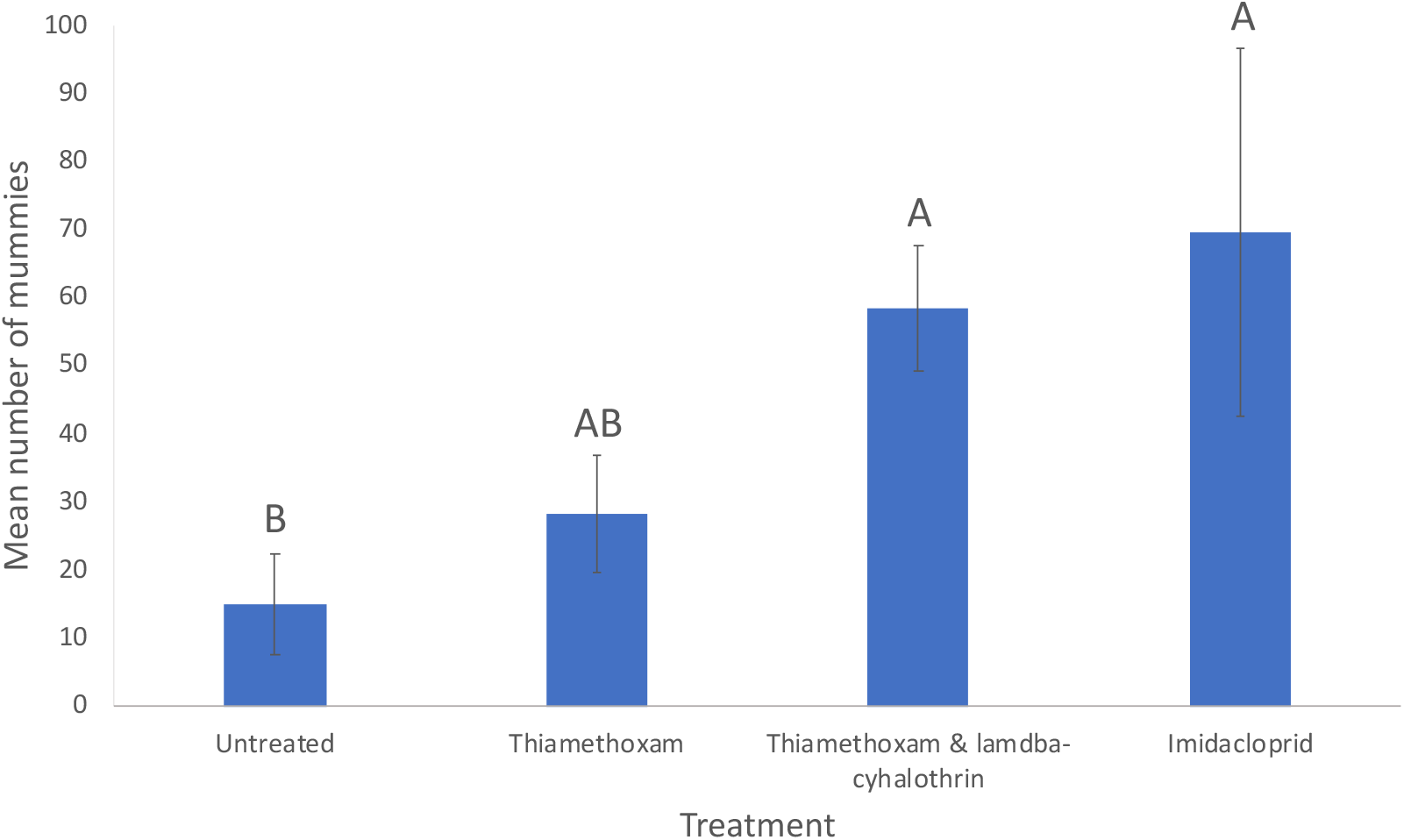
Average numbers of aphid mummies formed on untreated, and seed treated plants. Letters represent significant differences between treatments. Error bars represent standard errors of the mean.

The percentage of *A. colemani* reared from mummies (i.e., F1 parasitoids) was affected by treatment (F_(3,19)_=7.06, p<0.01), with a higher percentage of individuals reared on the thiamethoxam & lambda-cyhalothrin plants compared with the thiamethoxam and untreated plants (Fig. 6).

**Figure 6:**
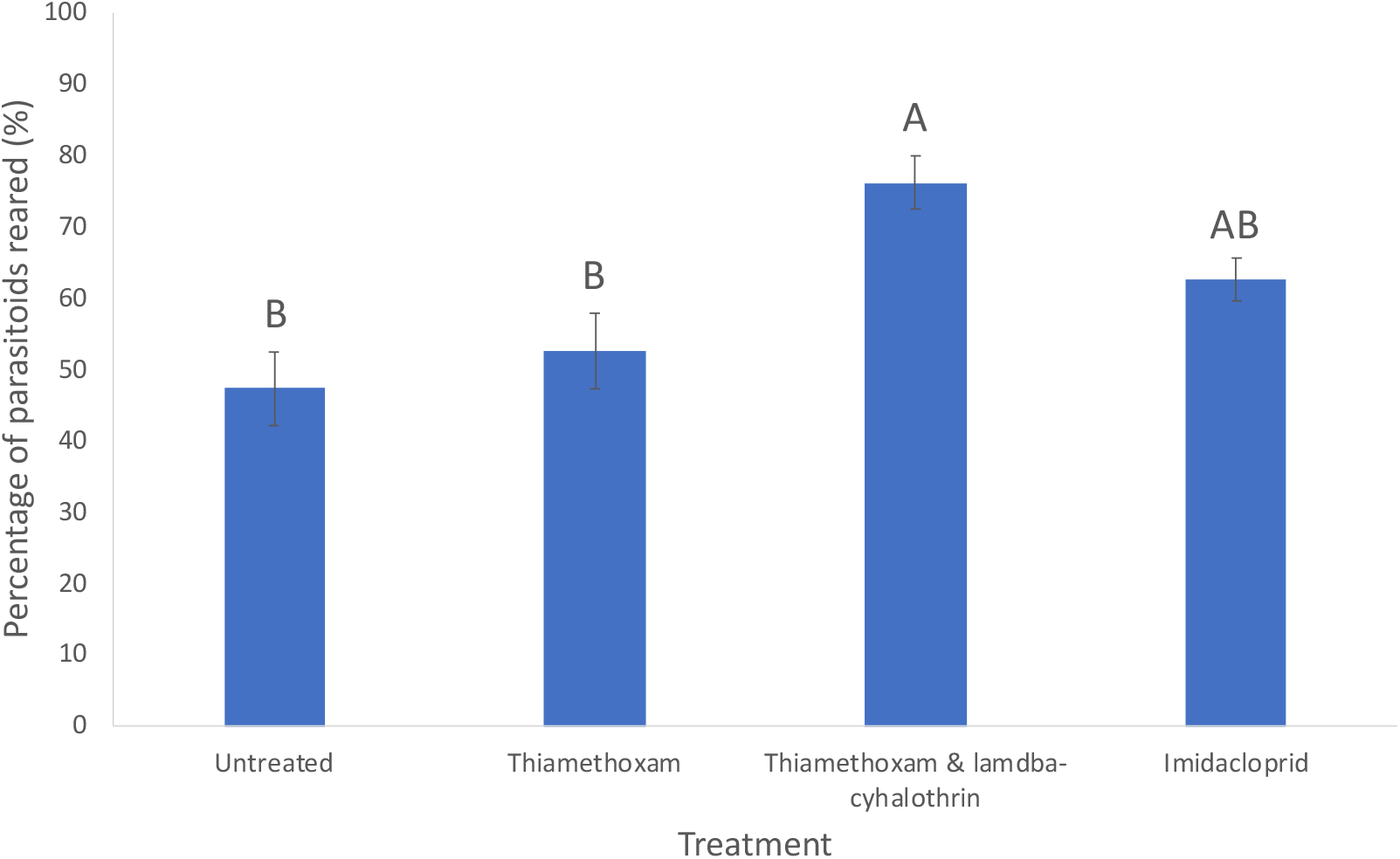
Average percentage of aphid mummies producing F1 *A. colemani* on untreated and seed treated plants. Letters represent significant differences between treatments. Error bars represent standard errors of the mean.

No significant treatment effect was identified for emergence, death rates or survival rates of F1 *A. colemani* collected as mummies from plants (Fig. S2).

#### 3.3.3. Trial 1a – Mummies collected from petri dishes

No significant difference among treatments was detected for the number of aphid mummies formed or the number of F1 *A. colemani* reared from mummies within the petri dishes (Fig. S3). Similar to F1 *A. colemani* reared from plants, the emergence rates and death rates of F1 *A. colemani* within the petri dishes were not significantly different between treatments (Fig. S4).

#### 3.3.4. Trial 1a – Overall measure of fitness of *A. colemani*

As an overall measure of fitness, the cumulative survival days were estimated for *A. colemani*. These were significantly different between treatments (ANOVA, F_(3,19)_=4.47, p<0.05). The fitness of *A. colemani* was higher in the thiamethoxam & lambda-cyhalothrin and Imidacloprid treatments, with both treatments significantly different to the untreated controls (Fig. 7), although this effect was no longer significant after Bonferroni correction.

**Figure 7:**
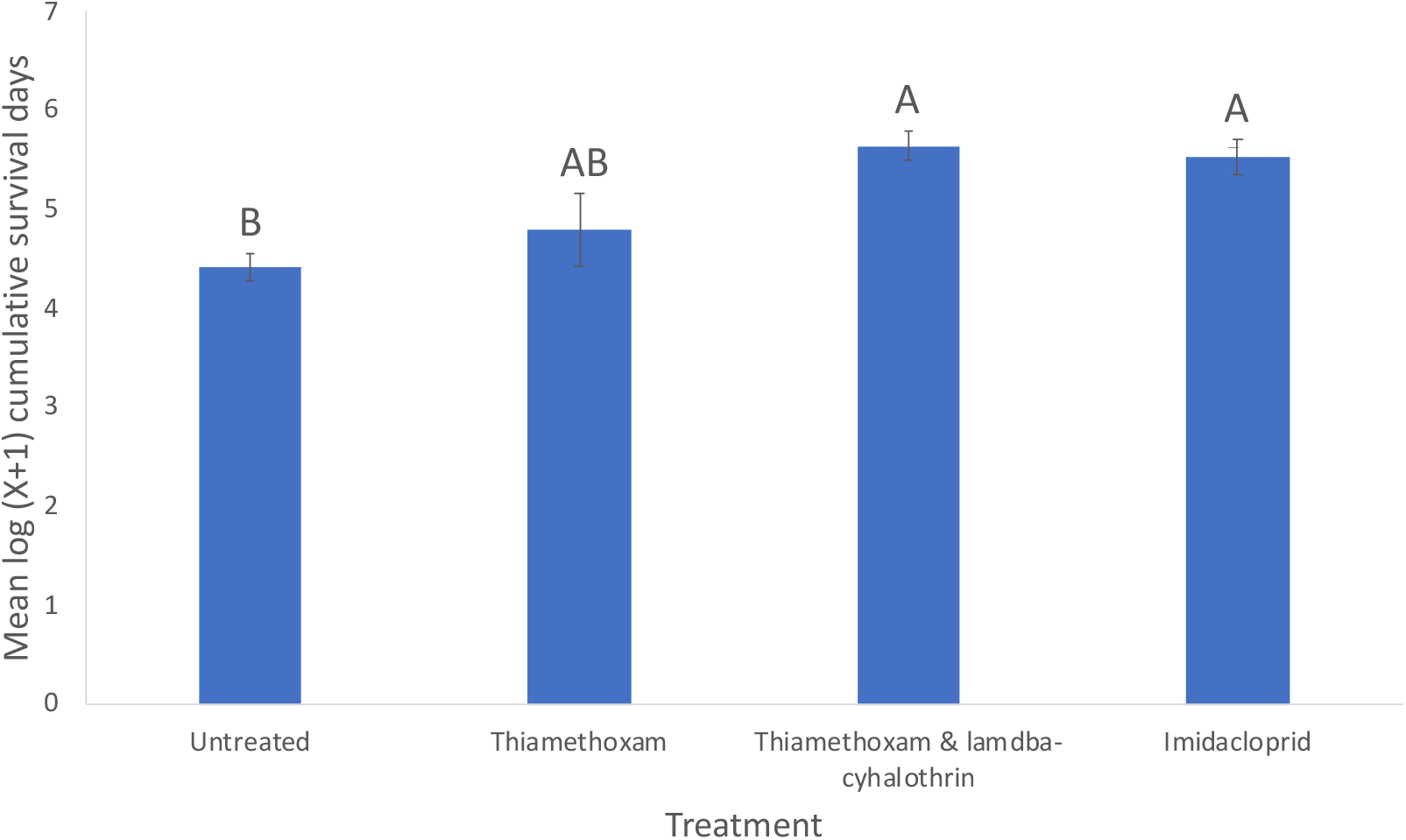
Total fitness measure for *A. colemani* on untreated and seed treated plants. Letters represent significant differences between treatments. Error bars represent standard errors of the mean.

### 3.4. Trial 1b – Behavioural experiments

When exposed to seed treated plants, there were no treatment effects for *M. persicae* behaviour. The period of time *M. persicae* spent inactive, walking, or showing any signs of movement did not differ across treatments (Fig. S5). When in the presence of *A. colemani*, there were also no treatment effects observed in *M. persicae* behaviour. There were no significant differences across treatments for the number of kicks when attacked by a parasitoid or the total number of kicks as a proportion of the number of attacks (Fig. S6).

When exposed to seed treated plants, there were some treatment effects on *A. colemani* behaviour. The duration of *A. colemani* antennal contact, resting, oriented walking, and the number of ovipositor contacts (A, B, and total), did not differ across treatments (Fig. S7). However, the duration of cleaning and searching by *A. colemani* did vary between treatments (cleaning, ANOVA, F_6,77_=5.02, p<0.001; searching, ANOVA, F_6,77_=5.41, p<0.001) (Fig. 8). When *A. colemani* were paired with aphids from the same treatments, *A. colemani* on imidacloprid-treated plants spent longer cleaning and less time searching than those from the other treatments, and those on thiamethoxam-treated plants the least time cleaning and the most time searching. When *A. colemani* from untreated plants were paired with *M. persicae* from insecticide seed treated plants, those paired with aphids from the thiamethoxam & lambda-cyhalothrin treatment spent the most time cleaning and the least time searching, and those paired with aphids from imidacloprid-treated plants the least time cleaning and the most time searching.

**Figure 8:**
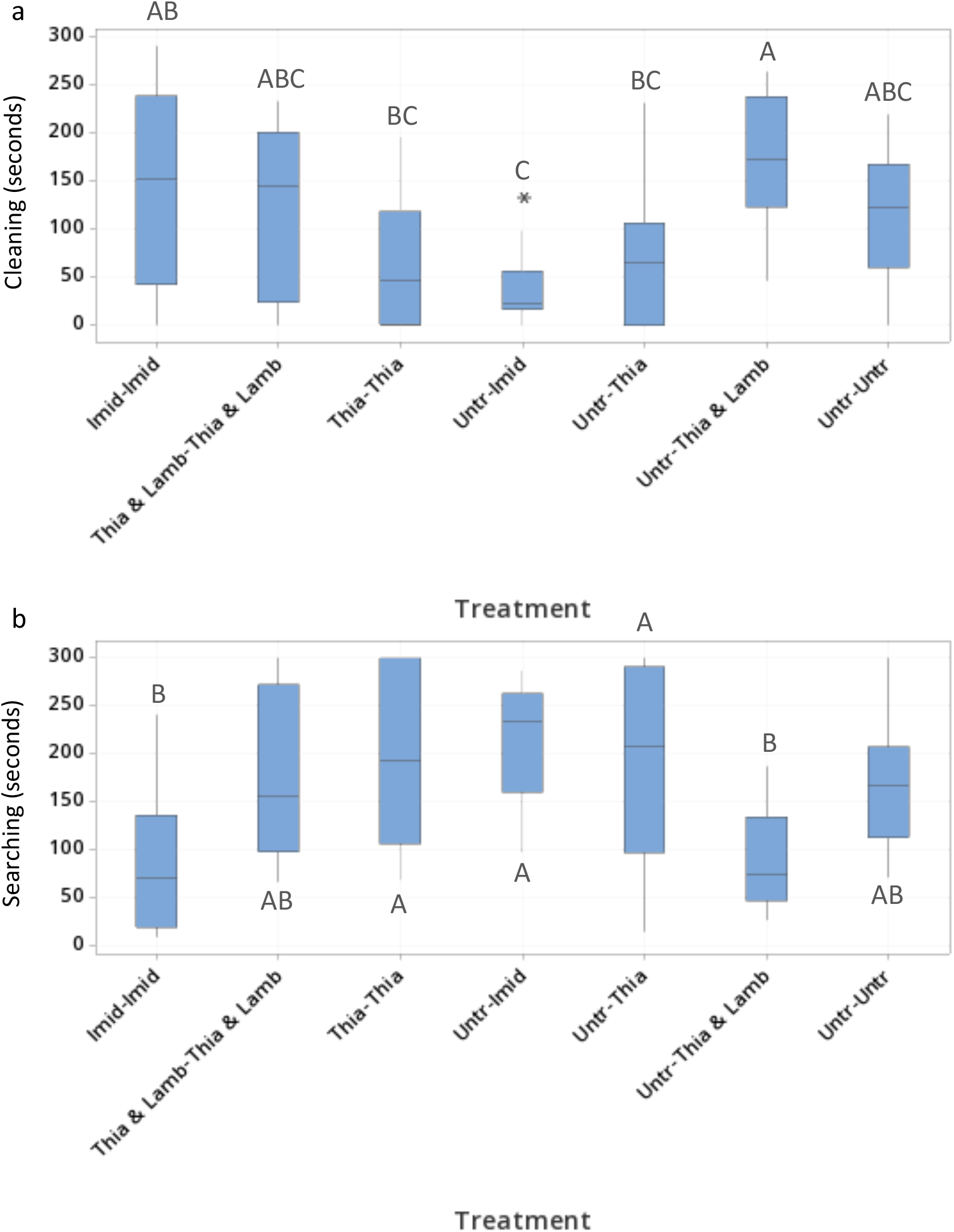
Treatment effects on time spent by *A. colemani* (a) cleaning and (b) searching. Letters represent significant differences between treatments. [‘Treatment’ indicates first the parasitoid treatment and second the paired *M. persicae* treatment; ‘Imid’ = Imidacloprid, ‘Thia & Lamb’ = Thiamethoxam & lambda-cyhalothrin, ‘Thia’ = Thiamethoxam, ‘Untr’ = Untreated. Outlier is shown as asterisk].

### 3.5. Mallada signatus

#### 3.5.1. Trial 2a – Initial mortality

No significant differences were found between treatments for each of the variables tested against *M. signatus* in Trial 2a. This included the initial mortality of 1^st^ stage lacewing larvae, 3^rd^ stage lacewing larvae, and total lacewings (Fig. S8). However initial mortality was higher for the 1^st^ stage larvae than the 3^rd^ stage larvae when treatments are paired (t(3)=5.42, p<0.05), although this effect was no longer significant after Bonferroni correction.

#### 3.5.2. Trial 2a – Pupation

There were no treatment effects for the number of 1^st^ stage larvae, 3^rd^ stage larvae, or total lacewings pupating, and there was no difference in the number of pupations between the 1^st^ stage and 3^rd^ stage larvae (Fig. S8). The number of days taken for the 1^st^ stage larvae to pupate after removal from canola plants averaged 10.57 days (standard deviation of 1.83 days), and this did not differ between treatments. The number of days taken for the 3^rd^ stage larvae to pupate after removal from the plants averaged 3.82 days (with a standard deviation of 1.00 days), and also did not differ between treatments. The number of days taken for the 1^st^ stage larvae to pupate was higher than for the 3^rd^ stage larvae (paired t(3)=115.39, p <0.001), as expected, due to their younger age.

The number of adults emerging did not vary between treatments for the 1^st^ stage larvae, 3^rd^ stage larvae or total lacewing larvae (Fig. S8). 1^st^ stage larvae had a mean total pupation success of 45% and the 3^rd^ stage larvae had a mean total pupation success of 61%. These were significantly different to one another (paired t(3)=-4.70, p<0.05), although this effect was no longer significant after Bonferroni correction. Furthermore, the number of days taken for the 1^st^ stage larvae, 3^rd^ stage larvae, and total lacewing larvae to emerge from the pupae did not differ between treatments (Fig. S9).

#### 3.5.3. Trial 2a – Longevity

There was no significant difference in adult longevity of *M. signatus* (from pupal emergence to death) between treatments (ANOVA, F_(3,58)_=0.25, p=0.862) (Fig. 9).

**Figure 9:**
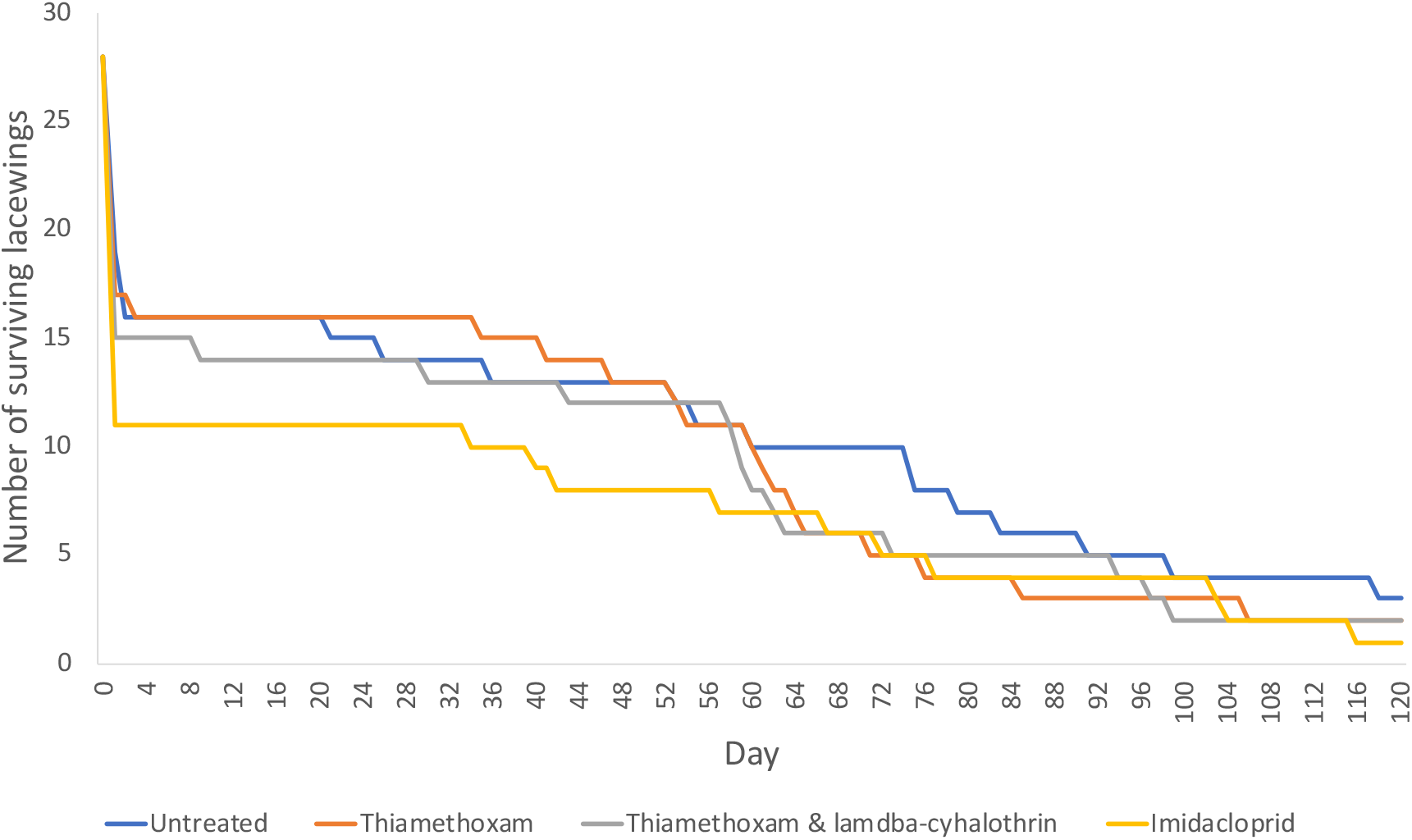
*M. signatus* survival when exposed to *M. persicae* on different insecticide seed treated canola plants.

#### 3.5.4. Trial 2b – Pupation

After 96 hours, only two lacewings had died, one in the untreated and one in the imidacloprid treatment. There were no significant differences between treatments in the number of lacewings pupating, or the number of days taken for *M. signatus* to pupate (Fig. S10).

## 4. Discussion

A large number of studies have shown insecticides to be toxic to parasitoids and predators. When natural enemies were exposed through tri-trophic interactions to insecticide seed treatments, Gontijo et al. (2018) found thiamethoxam caused mortality of the stink bug *Podisus nigrispinus* (Dallas), a predator of the fall armyworm *Spodoptera frugiperda* (Smith). Naveed et al. (2010) found lower field parasitism of the whitefly *Bemisia tabaci* (Gennadius) by aphelinid parasitoids when cotton seeds were treated with thiamethoxam and imidacloprid. Douglas et al. (2015) found although the pest slug *Deroceras reticulatum* (Müller) was unaffected by thiamethoxam, the chemical was passed to the predatory beetle *Chlaenius tricolor* Bonelli, which fed on the slugs, impairing or killing >60% of the beetles. We therefore expected the seed treatments tested here to have negative effects on *A. colemani* and *M. signatus*, both important beneficial insects in Australian canola, yet this study has not found strong evidence for these toxic effects. Through the behavioural studies undertaken with *M. persicae*, there were no treatment effects on aphid activity, which could be explained by the lack of detection of chemicals within the aphids themselves. Why then did we find significant differences between the seed treatments for some measures in *A. colemani*?

### 4.1. A. colemani

Aphidiines feed on honeydew produced by aphids (Wäckers, 2005). This could act as a pathway for chemicals to be transferred to the parasitoids via ingestion. There are records of imidacloprid detection within the honeydew of the striped pine scale *Toumeyella pini* (King) (Quesada et al., 2020), glucosinate sinigrin detection within the honeydew of *M. persicae* (Merritt, 1996), and terpenoid detection within the honeydew of *A. gossypii* (Hagenbucher et al., 2014). Perhaps the negligible levels of insecticide actives and their metabolites detected within aphids was caused by the excretion of these chemicals via their honeydew. Furthermore, imidacloprid has been shown to depress the excretion of honeydew by *M. persicae* by almost 95% within 24 hours (Nauen, 1995). In our study, parental *A. colemani* mummification and rearing success might have been affected by the seed treatments through honeydew ingestion, either by the parasitoid ingesting more of the chemical or producing less honeydew.

There were differences in the number of aphid mummies produced and the success of parasitoid emergence from mummies between the thiamethoxam and the thiamethoxam & lambda-cyhalothrin treatments. Although thiamethoxam was present at the same rate within both the seed treatments, the mass spectrometry results suggest a lower uptake of thiamethoxam in plants that were treated with lambda-cyhalothrin. *Aphidius colemani* exposed to the thiamethoxam & lambda-cyhalothrin treatment had a higher fitness than those exposed to the thiamethoxam only treatment. The presence of lambda-cyhalothrin could interact metabolically within the plant and/or aphid, causing lower levels of toxins to reach the beneficial organism. For example, in cotton leaves, the half-life of thiamethoxam is 1.9 days when present by itself but is reduced to 1.6 days when it is combined with lambda-cyhalothrin (Xuyang et al., 2013). Lambda-cyhalothrin could not be tested by mass spectrometry here, and therefore the presence and concentration of this chemical within the aphids is unknown.

So why are we seeing an apparent benefit of some insecticide seed treatments on *A. colemani* rather than negative effects as predicted? We are aware from the mass spectrometry results that the chemicals were taken up by the canola seedlings, as expected, although levels were higher than reported in other studies (e.g., listed by Krischik et al. (2015)). One explanation could be explained by a change in aphid behaviour, with the aphid becoming less fit and therefore less able to defend itself, for example (Booth et al. 2007). However, we did not see any differences in *M. persicae* behaviour after exposure to seed treatments, and so this explanation is unlikely. There are many studies that have shown hormoligosis in insects, a phenomenon which predicts that sub-harmful levels of an insecticide will be stimulatory to an organism through the provision of increased efficiency and increased sensitivity to respond to environmental changes (Luckey, 1968). Cutler (2013) lists a number of studies within which increased fecundity, stimulated oviposition, and decreased pupal mortality have been reported to occur in invertebrates. One such study found that DDT stimulated oviposition in a braconid parasitoid (Grosch and Valcovic, 1967). This could be a plausible explanation and matches with the mass spectrometry results which demonstrate only low quantities of the chemicals reaching the aphids (and thus reaching *A. colemani*).

We found *A. colemani* from the imidacloprid treatment spent the shortest time (and those from the thiamethoxam treatment the longest time) searching for same-treatment hosts than parasitoids within the other treatments. Conversely, when *A. colemani* from the untreated plants were provided with *M. persicae* from the chemical seed treatments, those paired with the imidacloprid and thiamethoxam treatments spent more time searching than for the other treatments. This could suggest thiamethoxam inhibited the ability of *A. colemani* to locate *M. persicae*, a phenomenon recorded by Mustard et al. (2020), who determined this insecticide directly affected the olfactory perception of odours and the foraging ability of honeybees (*Apis mellifera* L.). Walking speed was not calculated for each seed treatment, however, may have been affected by thiamethoxam, such as in the case of the spotted lady beetle *Coleomegilla maculata* DeGeer (Bredeson et al. 2015).

Significant effects of seed treatments were constrained to the F1 *A. colemani* produced on the insecticide seed treated plants. This suggests that the seed treatment effects on *A. colemani* may not be long-lasting. This is congruent with other insect studies. For example, Nauen (1995) found that within 24 hours of being removed from imidacloprid-treated leaves, *M. persicae* reversed their immediate behavioural responses and begun increasing in weight and producing more honeydew.

### 4.2. M. signatus

We found no significant effects of insecticide seed treatments on *M. signatus* mortality, larval and pupal survival, larval and pupal duration, or adult longevity. Other studies have found lacewings to be tolerant to a range of agricultural insecticides. For example, imidacloprid was shown to have low toxicity to the green lacewing *Chrysoperla rufilabris* (Burmeister), causing 1-11% mortality (Mizell III and Sconyers, 1992). A field study in sorghum crops also found imidacloprid seed treatments to have little/no impact on lacewings, but negatively affect other predators such as the ladybird beetle *Hippodamia convergens* Guerin (Krauter et al., 2001). Directly comparing a parasitoid, Sterk et al. (1999) examined the mortality of the green lacewing *Chrysoperla carnea* and *Aphidius matricariae* Haliday after exposure to lambda-cyhalothrin, ranking the chemical as ‘harmful’ to *A. matricariae* (causing >75% mortality), yet ‘harmless’ to *C. carnea* (causing <25% mortality).

Conversely, Gontijo et al. (2014) explored the impacts of thiamethoxam-treated sunflower seeds on the green lacewing *C. carnea* and found thiamethoxam was toxic, reducing the fecundity and survival of adults. *Chrysoperla* sp. adults were also reduced in numbers in soybean fields grown from thiamethoxam-treated seed (Seagraves and Lundgren, 2012). Gontijo et al. (2014) suggest the greater impact of seed treatments on adult lacewings may be, in part, due to their greater consumption of extra-floral nectar. Thus, the lack of significant effects in our study could be due to a lack of extrafloral or floral nectar feeding, given the canola plants used here were not at the flowering stage. Juveniles were fed contaminated aphids, yet after pupation, adults were fed untainted pollen.

### 4.3. Conclusion

It is possible the initial exposure time to chemicals in our study may have been insufficient to exhibit the full lethal or sublethal effects against *M. signatus* and *A. colemani*. Mortality of *A. colemani* when exposed to the volatile oil UDA-245 was found to be significantly greater at 48 hours than at 24 hours exposure, through contact toxicity, but not through residual toxicity (Bostanian et al. 2005). A similar finding has been observed for the aphidiine, *L. testaceipes;* after exposure to azadirachtin, survival was found to significantly reduce from 69-80% at 24 hours after treatment to 28-33% at 48 hours after treatment (Tang et al. 2002). In their study, Anjum & Wright (2016) showed the intrinsic toxicity of lambda-cyhalothrin was greater against *M. persicae* when exposure time increased from 24 hours to 120 hours. These comparative studies suggest the results we observed could be quite different if the exposure time of aphids (and/or natural enemies) was extended. This is something that could be investigated through further experimentation.

Selectivity of insecticides to beneficial arthropods is important for the implementation of IPM programs and conservation biological control (Bacci et al., 2009, Jansen et al., 2008, Sterk et al., 1999). Surprisingly, we saw little evidence of negative toxic effects against *A. colemani* and *M. signatus* when exposed to aphids that had fed on insecticide seed treated canola seedlings. It is important these trials are repeated using semi-field or field studies, given chemical impacts can vary considerably with laboratory trials such as those conducted here (Hassan et al., 1988). Further experiments should also be undertaken with other species to determine how widespread these patterns are across taxa. One species of particular importance in grain crops is *Diaeretiella rapae* (M’Intosh), which is the most commonly found aphidiine in canola fields in Australia (Ward et al., 2021). Globally, there is very little data on the toxicity of insecticide products against this species. IPM programs should take into account the impact of insecticide seed treatments on both parasitoids and predators, and in particular sublethal effects, which can be easily overlooked, but which can have considerable impact on ecosystem services (Gontijo et al., 2014).

## Acknowledgements

We would like to begin by thanking Biological Services for providing parasitoids for experimental work, Kelly Angel at the Birchip Cropping Group (BCG) for providing canola seed, David Landmeter at Syngenta and the staff at the Shepparton branch of Eurofins Agrisearch services for their assistance coating the canola seeds. We acknowledge Ina Smith from CSIRO and Richard O’Hair at the University of Melbourne for providing their time to discuss chemical testing, Toshifumi Nakao for his advice, and Andrew Warden, Greg Dojchinov and Matthew Taylor for their mass spectrometry work. We extend our gratitude to Alex Gill, for his assistance in maintaining colonies during the Covid-19 lockdown, Perran Ross for his microscope photography training, and Kathy Overton, for the provision of references. Thank you also to Sarina Macfadyen and Hazel Parry for their advice. This research was supported by a GRDC investment that seeks to deliver new knowledge to improve the timing of pest management decisions in grain crops to grain growers: CSE00059. This work was further supported by the Albert Shimmins Fund.

