## Supplementary material for "The effects of insecticide seed treatments on green peach aphid *Myzus persicae* (Sulzer) (Homoptera: Aphididae) parasitism by *Aphidius colemani* Viereck (Hymenoptera: Aphidiidae) and predation by *Mallada signatus* (Schneider) (Neuroptera: Chrysopidae)": Fig. S

*Table S1*: Modified aphid and parasitoid behavioural table from Bilodeau et al. (2013).

| ***Organism*** | ***Behaviour*** | ***Remarks*** |
| --- | --- | --- |
| *Aphid* | Inactivity | Aphid is completely immobile. |
|  | Walking/running | Aphid walks/runs. |
|  | Cornicle secretion | Aphid secretes from its cornicles. |
|  | Movement | Movement; not walking/running i.e., kicking without parasitoid present, twitching. |
|  | Resistance | Aphid being contacted by wasp actively resists/kicks (only relevant when parasitoids present). |
| *Parasitoid* | Antennal contact | Palpation of aphid with antennae. |
|  | Non-oriented contact | Apparently accidental contact with aphid. |
|  | Resting | Parasitoid is completely immobile. |
|  | Oriented walking | Walking toward aphid or reaching aphid. |
|  | Cleaning | Cleaning itself with legs and/or mouthparts. |
|  | Ovipositor contact A | Parasitoid strikes aphid with ovipositor on abdomen or thorax. |
|  | Ovipositor contact B | Parasitoid strikes aphid with ovipositor on head or appendages. |
|  | Searching | Walking rapidly or flying apparently to disperse. |

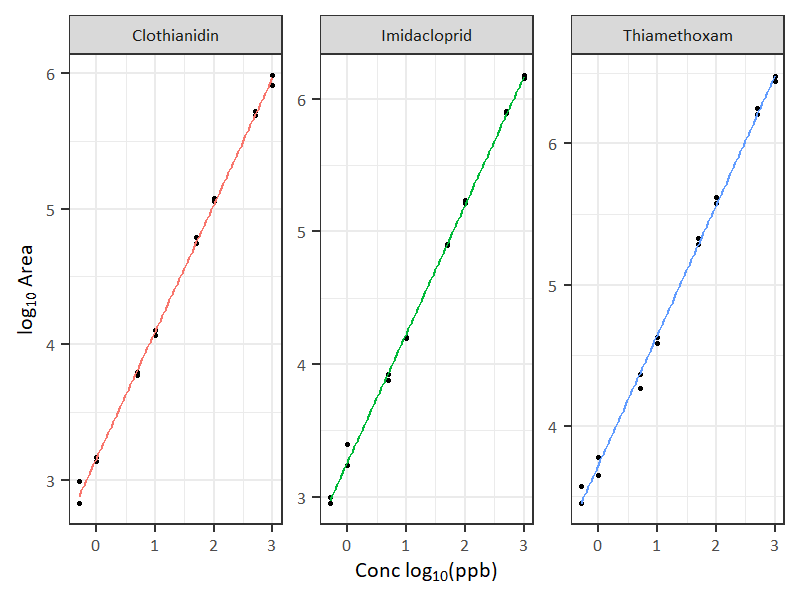

*Figure S1:* Standard curves for imidacloprid, thiamethoxam and clothianidin, expressed in parts per billion.

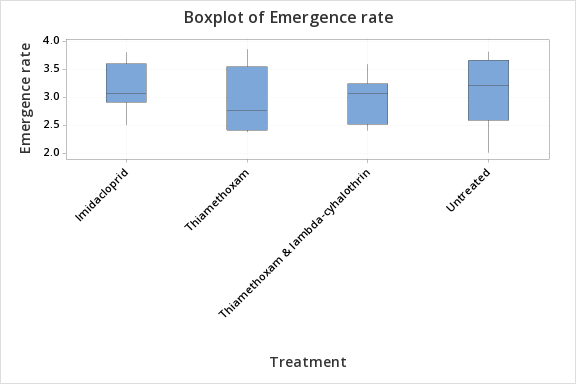

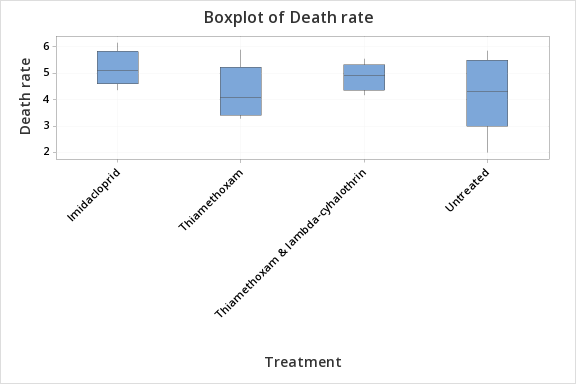

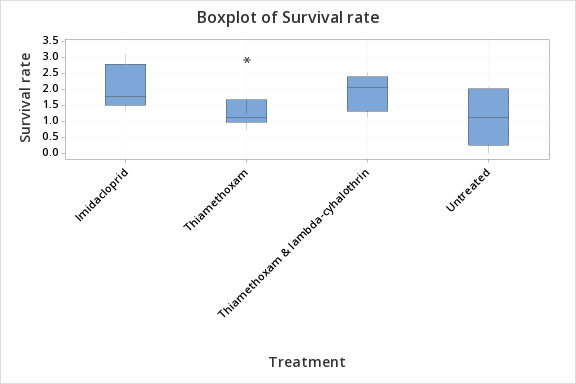

a

b

c

Survival rate (a-b)	Death rate (a)	Emergence rate (b)

*Figure S2:* Effect of seed treatments on a) emergence, b) death, and c) survival rates of F1 *A. colemani* on canola plants. No significant treatment effect was identified for emergence rates (ANOVA, F_(3,19)_=0.27, p=0.845), death rates (ANOVA, F_(3,19)_=1.28, p=0.311) or survival rates (ANOVA, F_(3,19)_=1.91, p=0.163) of F1 *A. colemani.* [Outlier is shown as an asterisk; Not to scale].

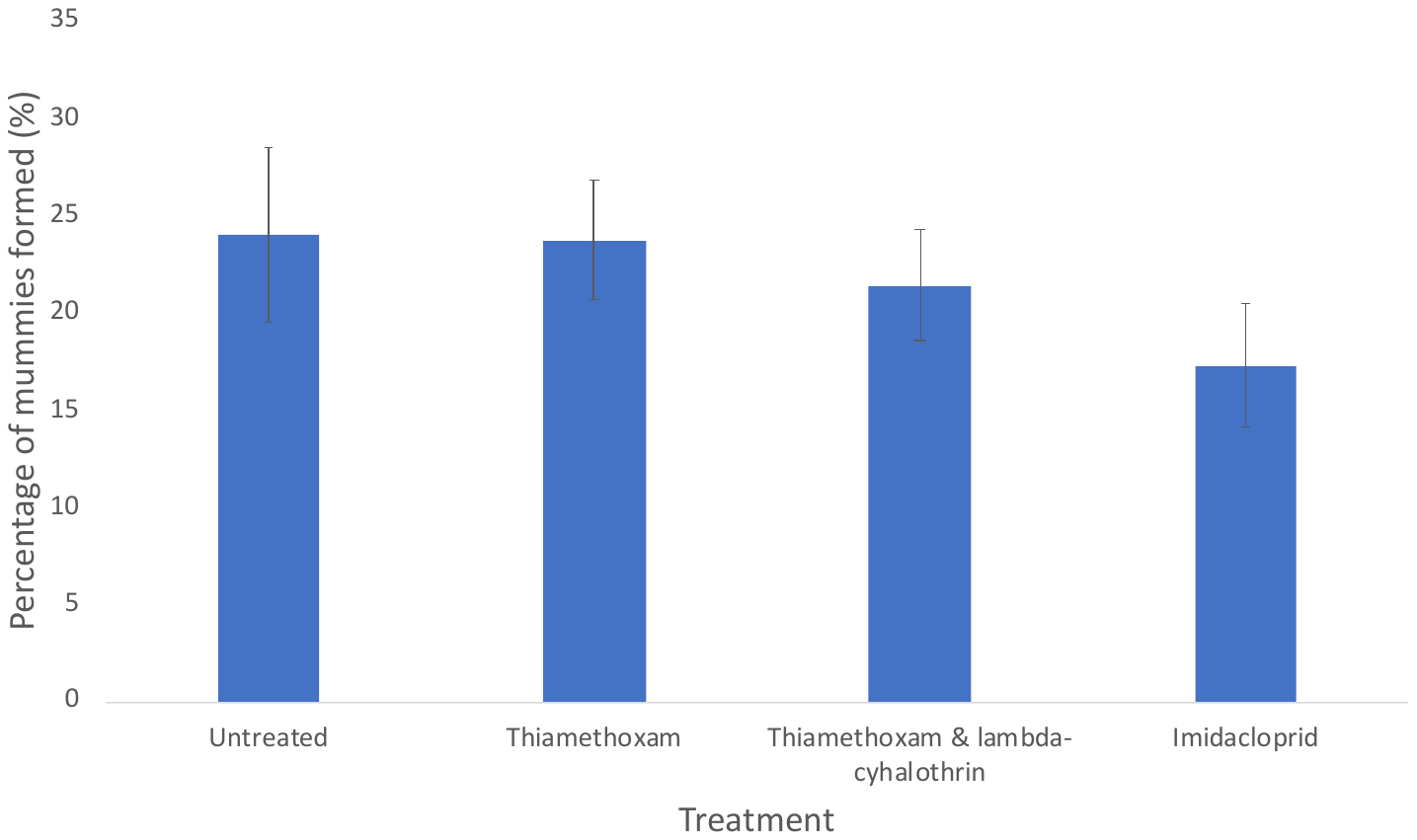

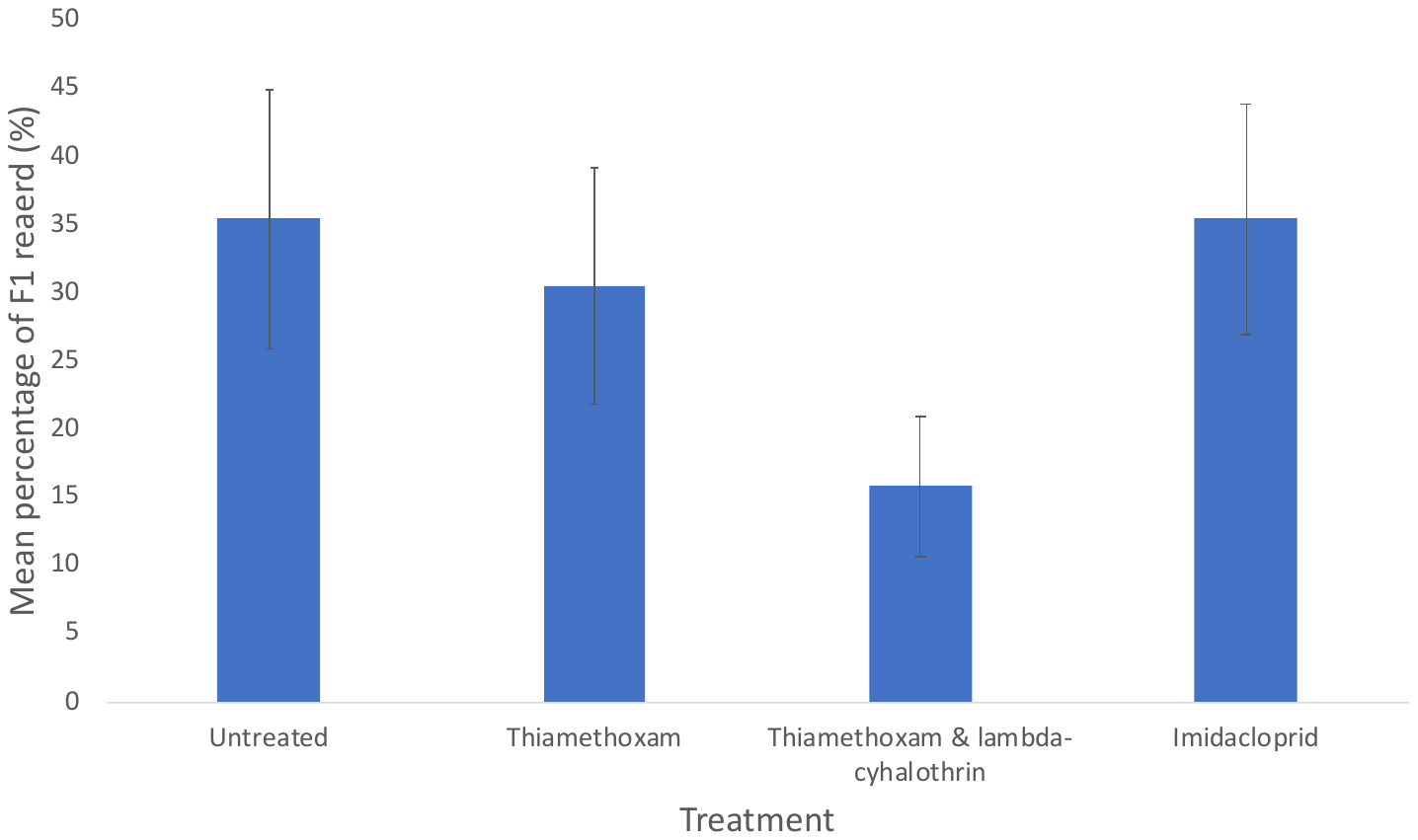

a

b

*Figure S3:* Effect of treatments on a) aphid mummies formed from total population and b) F1 *A. colemani* reared from mummies, within petri dishes. No significant difference among treatments was detected between the number of aphid mummies formed (ANOVA, F_(3,59)_=0.57, p=0.636), or the F1 *A. colemani* numbers reared from mummies (ANOVA, F_(3,20)_=1.13, p=0.360) within the petri dishes [Error bars are standard errors of the mean; Not to scale].

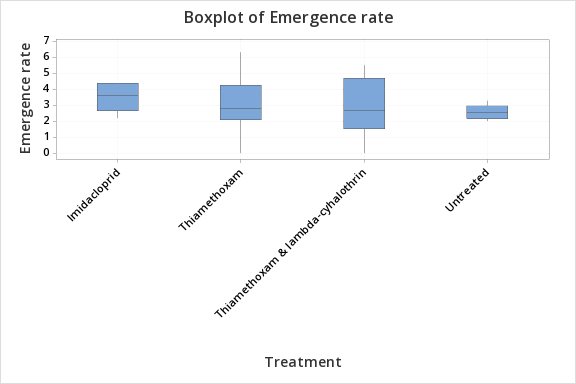

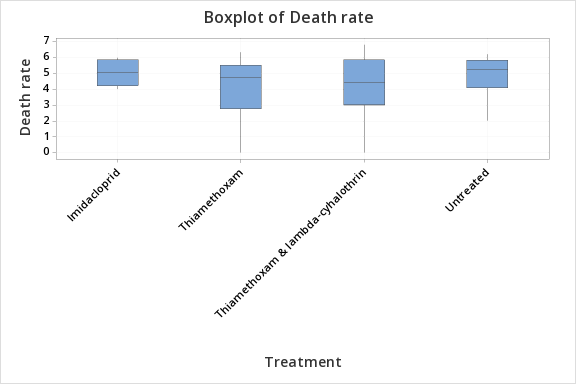

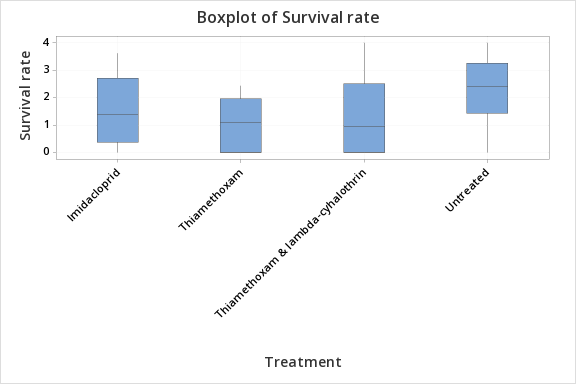

a

b

c

Survival rate (a-b)	Death rate (a)	Emergence rate (b)

*Figure S4:* Effect of treatments on a) emergence, b) death, and c) survival rates of F1 *A. colemani* in petri dishes. No significant treatment effect was identified for emergence rates (ANOVA, F_3,20_=0.38, p=0.768), death rates (ANOVA, F_3,20_=0.40, p=0.752). or survival rates (ANOVA, F_(3,20)_=0.93, p=0.445) of F1 *A. colemani.* [Not to scale].

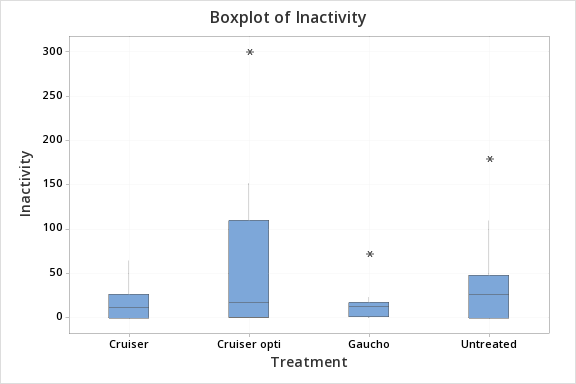

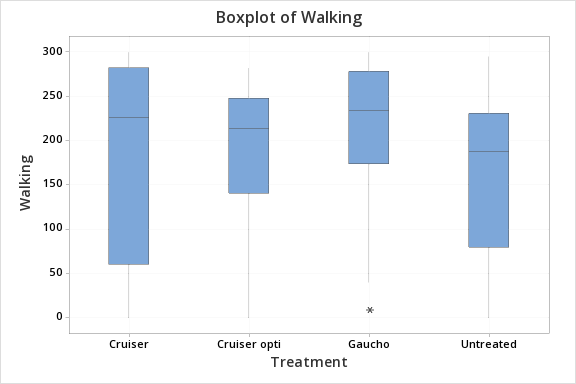

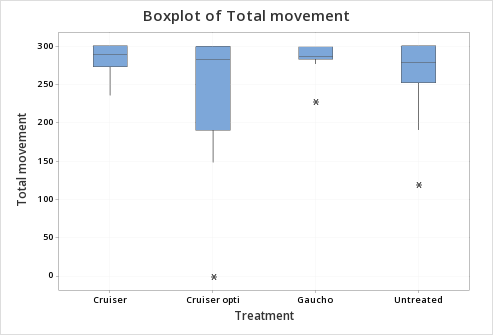

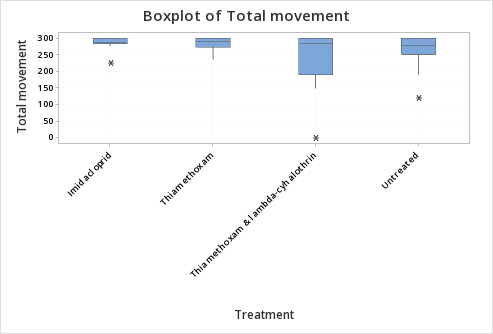

a

b

c

Total movement (seconds)	Walking (seconds)	Inactivity (seconds)

Imidacloprid

Thiamethoxam

Thiamethoxam & lambda-cyhalothrin

Untreated

Figure S5: Effect of treatments on duration of *M. persicae* a) inactivity, b) walking, and c) moving. The period of time *M. persicae* spent a) inactive, b) walking, or c) showing any signs of movement did not differ across treatments (inactivity, ANOVA, F_3,44_=1.88, p=0.147; walking, ANOVA, F_3,44_=0.59, p=0.624; total movement, ANOVA, F_3,44_=1.86, p=0.151). [Outliers are shown as asterisks].

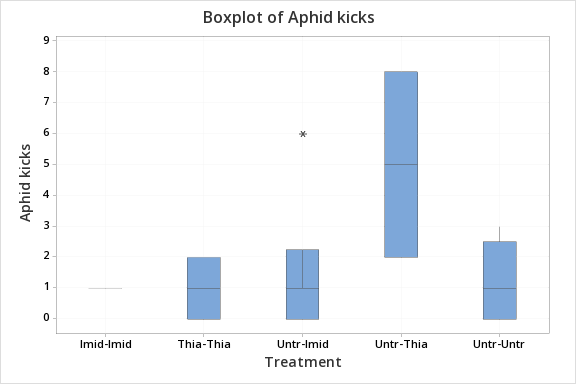

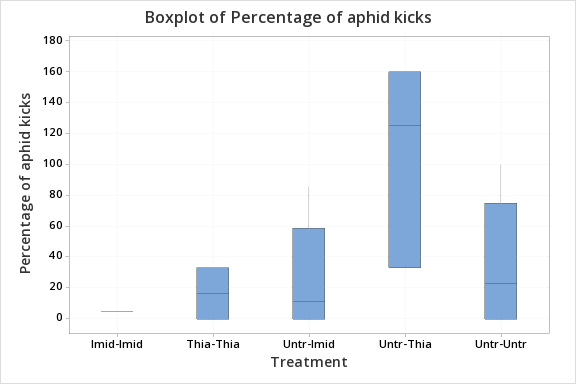

a

b

*Figure S6:* Effect of treatments on a) number of kicks by *M. persicae* and b) number of *M. persicae* kicks as a proportion of the number of *A. colemani* attacks. [Percentage is over 100% when aphids kick without being attacked]. There were no significant differences across treatments for the number of kicks when attacked by a parasitoid (ANOVA, F_4,12_=1.99, p=0.161) or the total number of kicks as a proportion of the number of attacks (ANOVA, F_4,12_=2.26, p=0.123) [‘Treatment’ indicates first the parasitoid treatment and second the paired *M. persicae* treatment; ‘Imid’ = Imidacloprid, ‘Thia & Lamb’ = Thiamethoxam & Lambda-cyhalothrin, ‘Thia’ = Thiamethoxam, ‘Untr’ = Untreated. Outlier is shown as an asterisk].

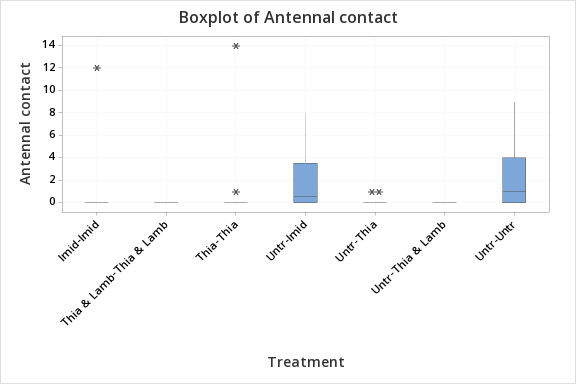

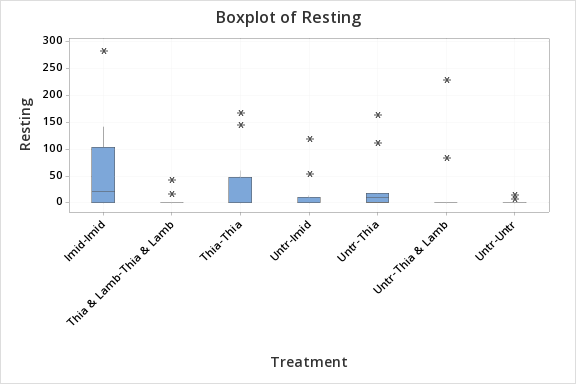

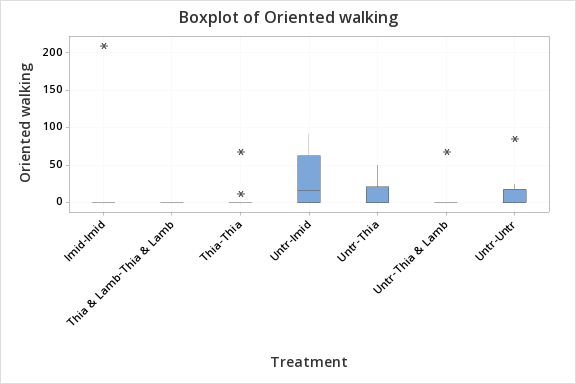

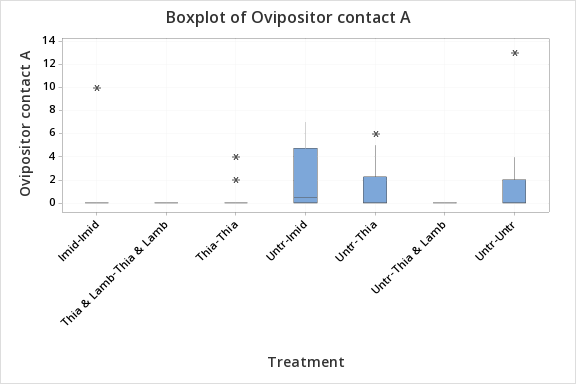

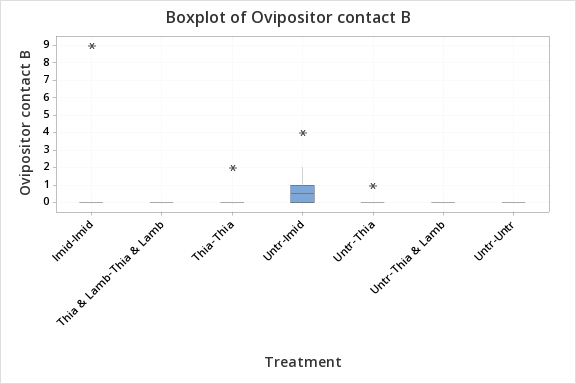

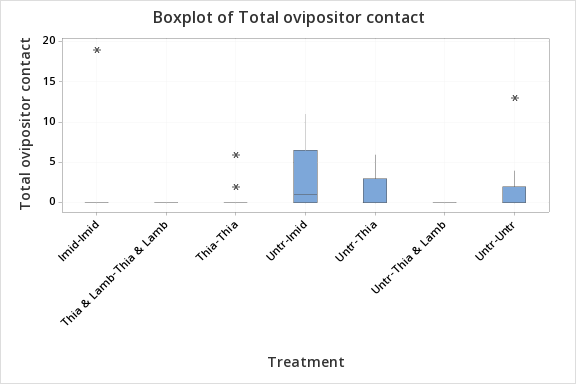

a	b

c	d

e	f

*Figure S7:* Effect of treatments on *A. colemani* behaviour. The duration of *A. colemani* a) antennal contact, b) resting, c) oriented walking, d) the number of A ovipositor contacts, e) the number of B ovipositor contacts, and f) the total number of ovipositor contacts, did not differ across treatments (antennal contact, ANOVA, F_6,77_=1.61, p=0.155; resting, ANOVA, F_6,77_=1.64, p=0.148; oriented walking, ANOVA, F_6,77_=1.15, p=0.340; ovipositor contact A, ANOVA, F_6,77_=1.75, p=0.120; ovipositor contact B, ANOVA, F_6,77_=1.32, p=0.257; ovipositor contact total, ANOVA, F_6,77_=1.52, p=0.183) [‘Treatment’ indicates first the parasitoid treatment and second the paired *M. persicae* treatment; ‘Imid’ = Imidacloprid, ‘Thia & Lamb’ = Thiamethoxam & Lambda-cyhalothrin, ‘Thia’ = Thiamethoxam, ‘Untr’ = Untreated. Outliers are shown as asterisks].

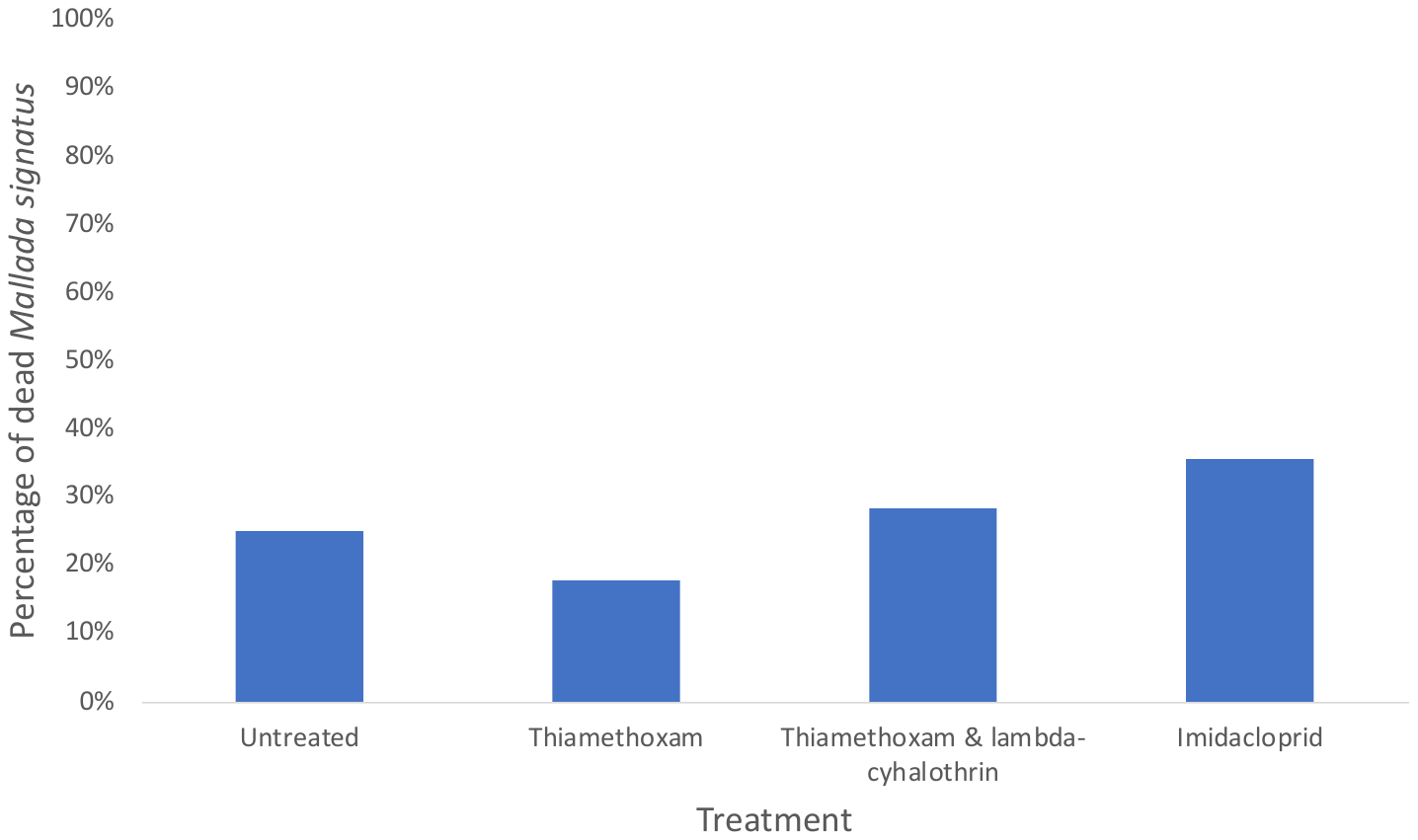

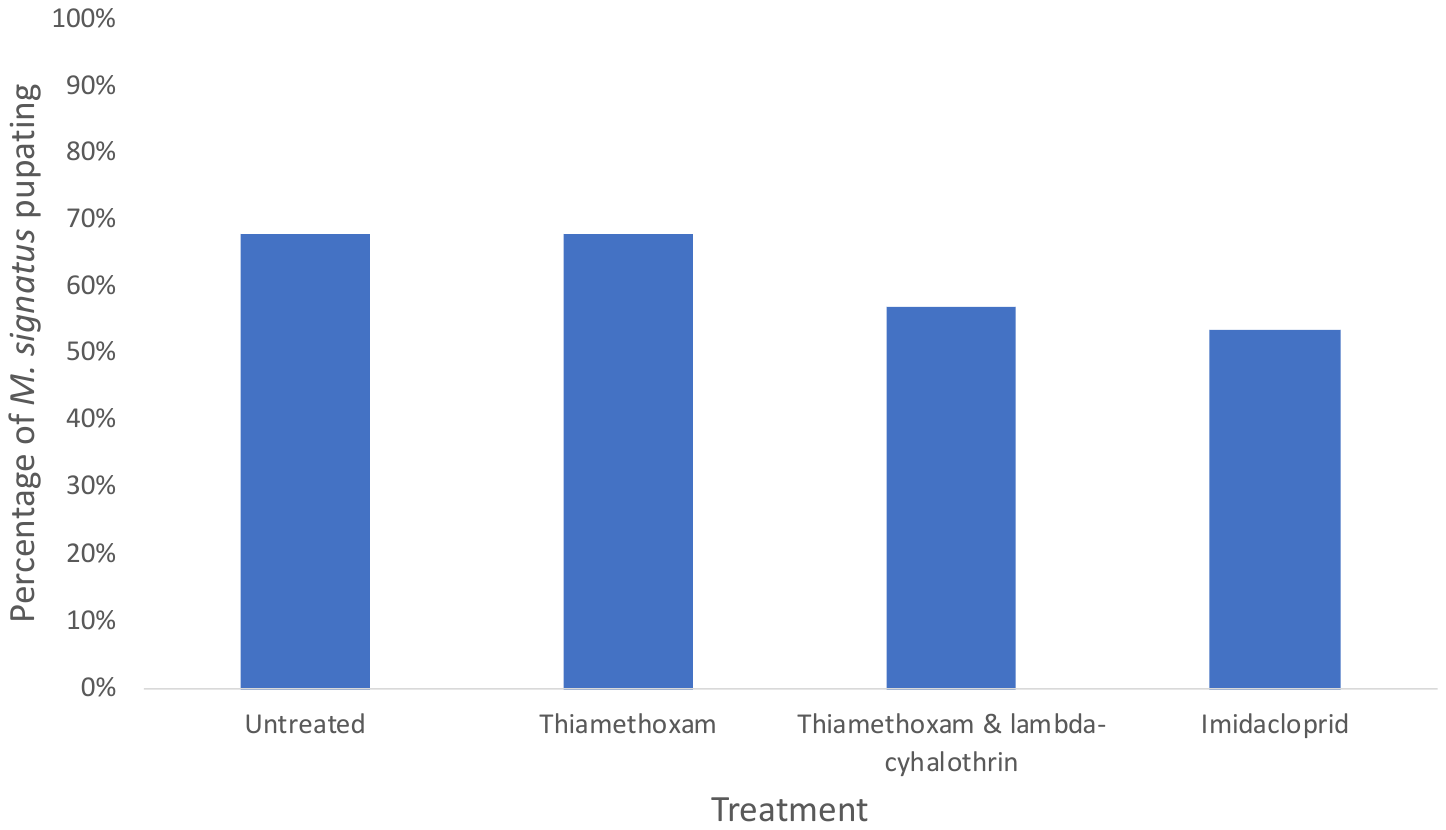

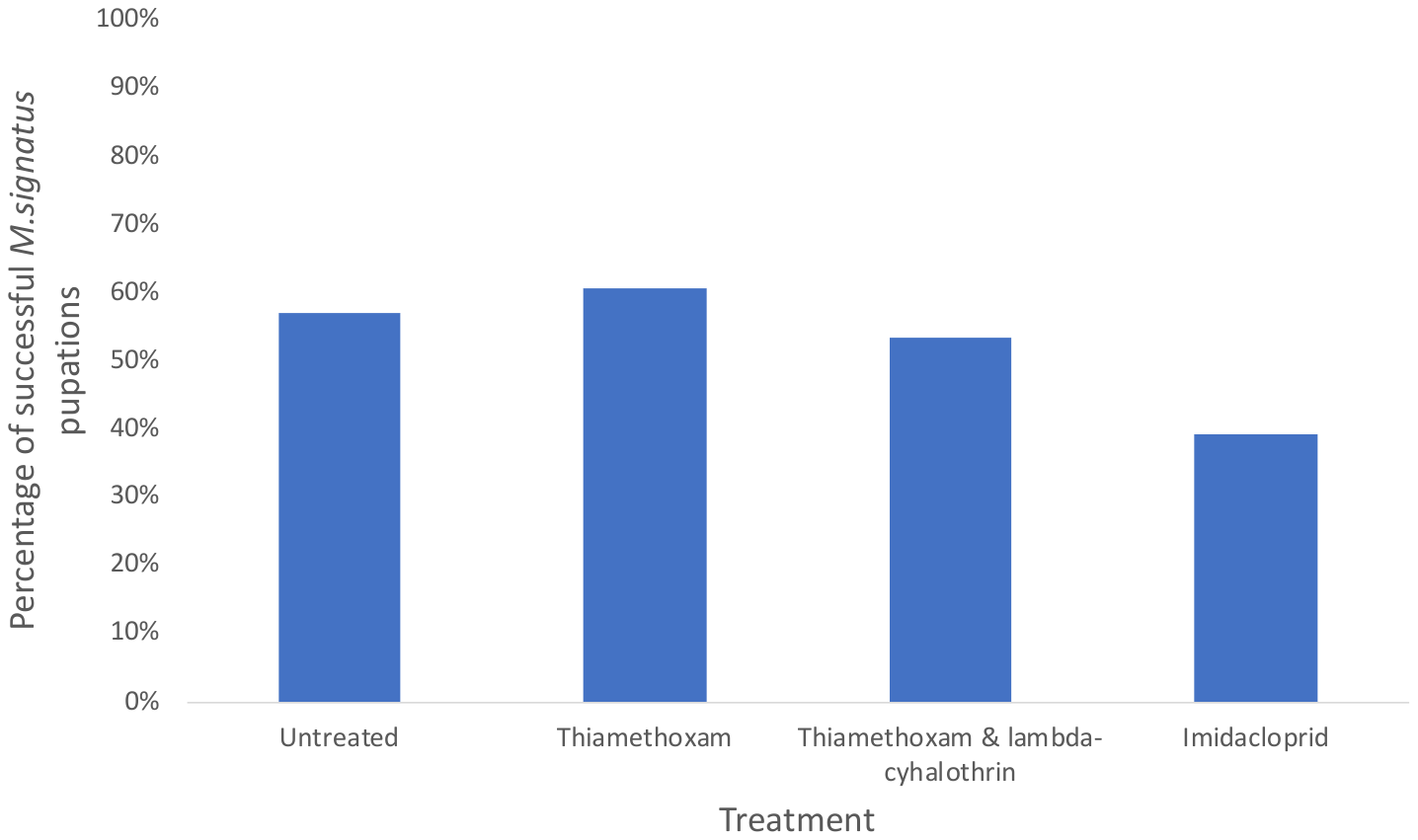

a

b

c

*Figure S8:* Effect of treatments on a) the initial mortality of total lacewings, b) the percentage of total lacewings pupating, and c) the number of successful pupations for total lacewings. There was no significant treatment effect on the initial mortality of 1^st^ stage lacewing larvae (contingency test, χ^2^=0.898, df=3, p=0.826, N=56), 3^rd^ stage lacewing larvae (χ^2^=3.500, df=3, p=0.321, N=56), and total lacewing larvae (χ^2^=2.367, df=3, p=0.500, N=112). There were also no significant treatment effects for the number of 1^st^ stage larvae (contingency test, χ^2^=1.436, df=3, p=0.697, N=56), 3^rd^ stage larvae (χ^2^=3.500, df=3, p=0.230, N=56), or total lacewings (χ^2^=1.925, df=3, p=0.588, N=112) pupating. The number of successful pupations did not significantly differ between treatments for the 1^st^ stage larvae (contingency test, χ^2^=0.795, df=3, p=0.851 N=56), 3^rd^ stage larvae (χ^2^=2.695, df=3, p=0.441, N=56), or total lacewing larvae (χ^2^=2.973, df=3, p=0.396, N=112).

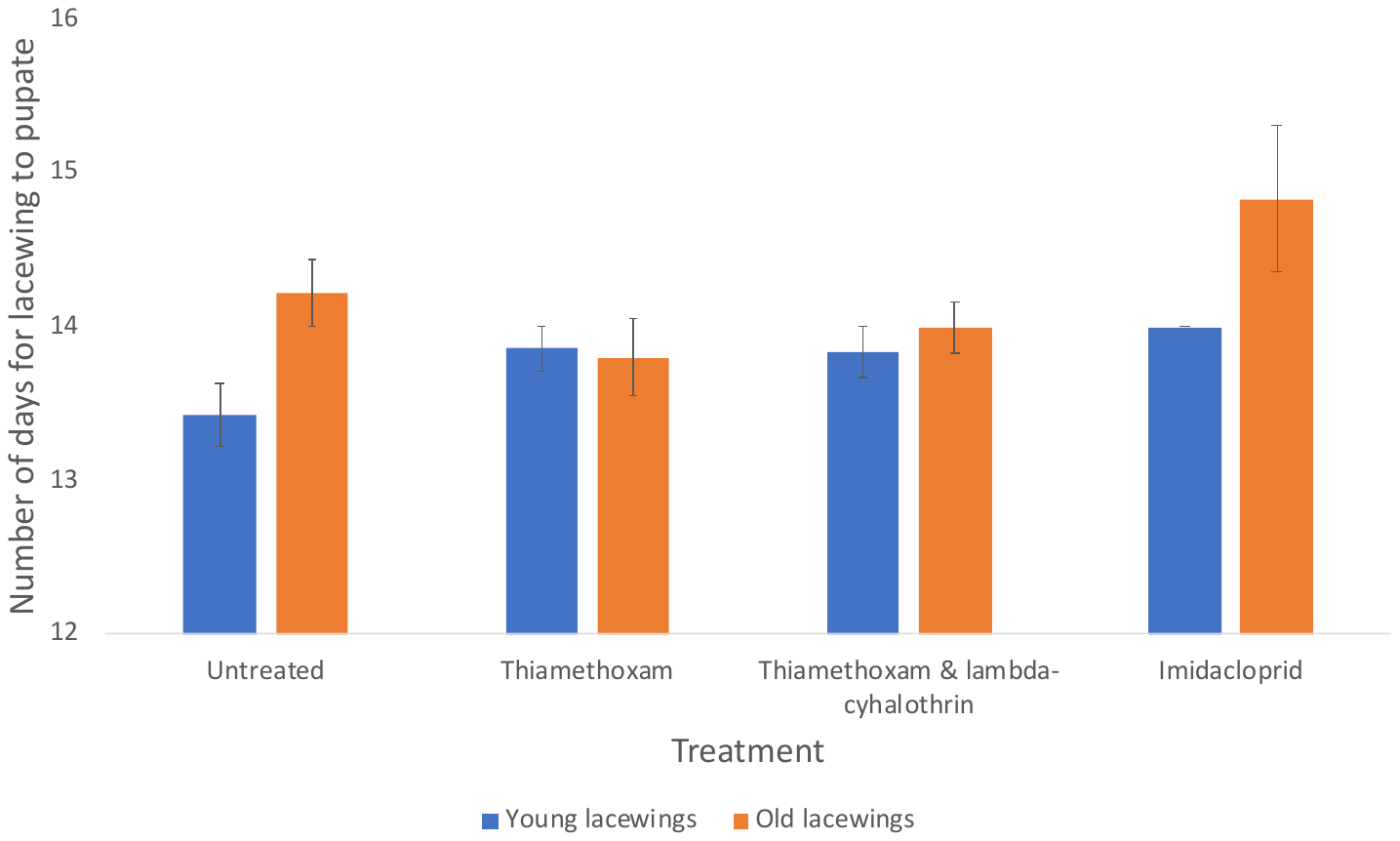

*Figure S9:* Effect of treatments on the number of days for 1^st^ stage and 3^rd^ stage larvae lacewings to emerge from pupae. The number of days taken for the total adult lacewings to emerge from the pupae did not differ between treatments (ANOVA, F_3,55_=2.20, p=0.099), which was also the case when considering 1^st^ stage larvae (ANOVA, F_3,21_=2.38, p=0.099) and 3^rd^ stage larvae (ANOVA, F_3,30_=2.38, p=0.090) separately.

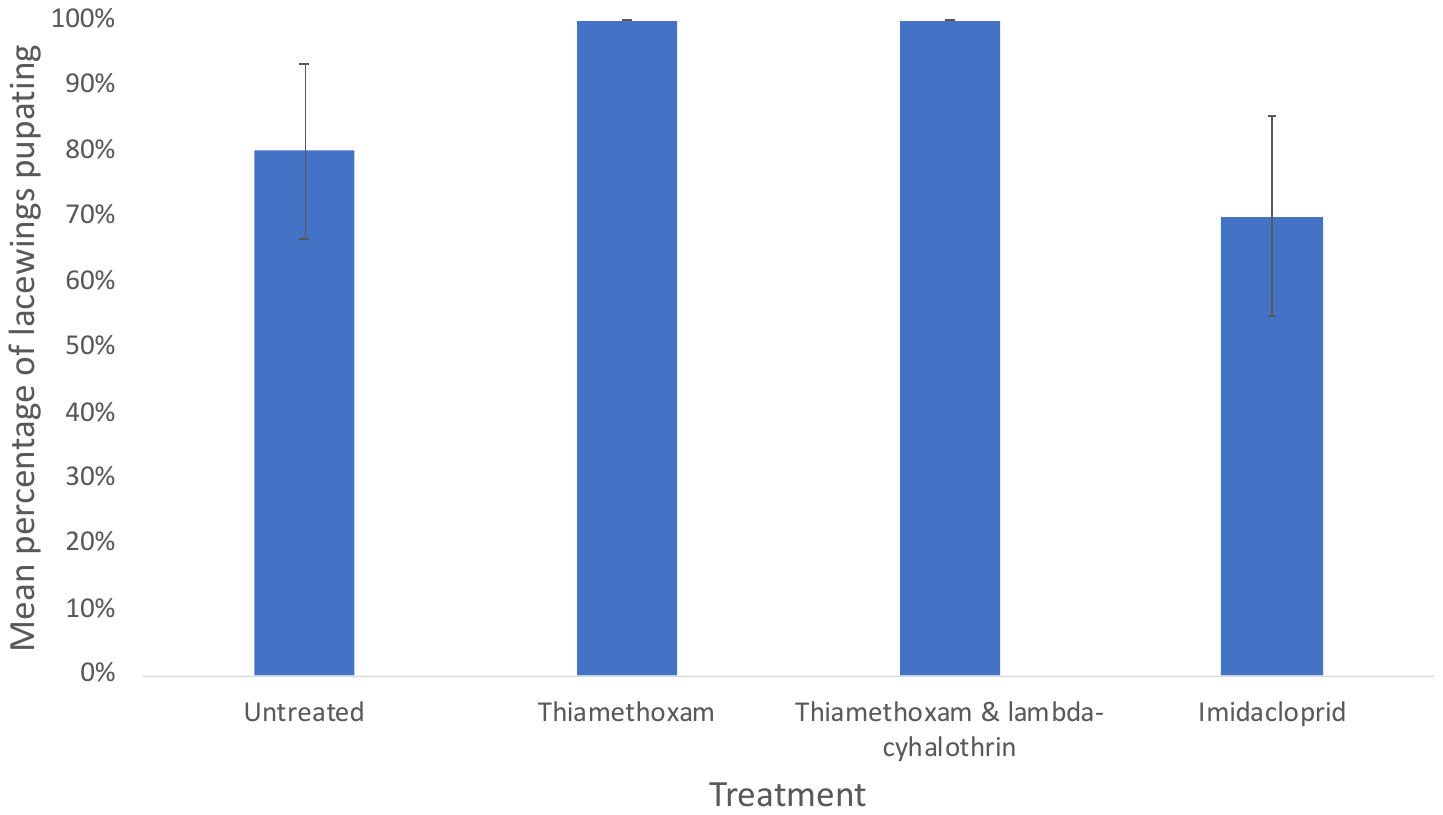

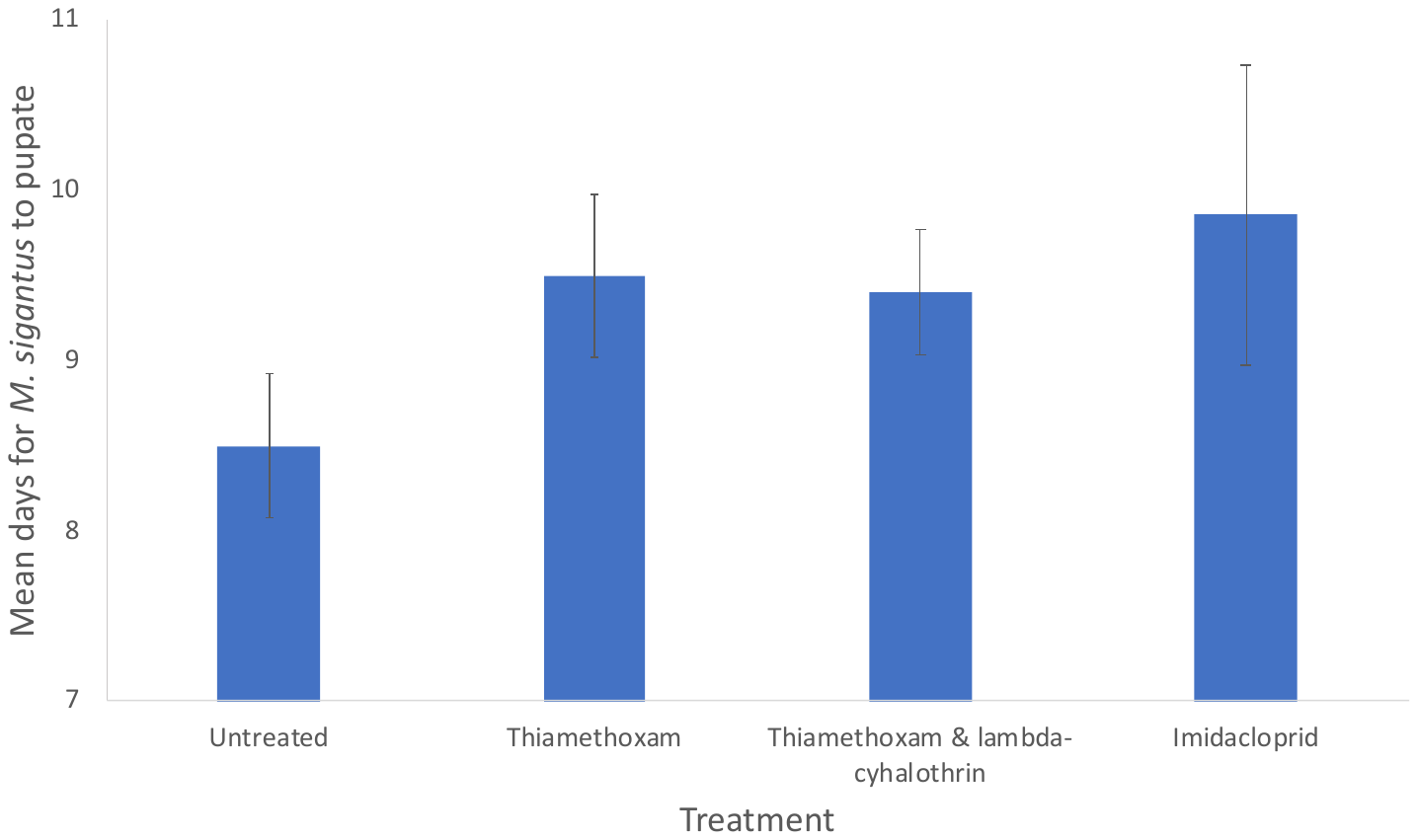

a

b

*Figure S10:* Effect of treatments on a) the mean percentage of *M. signatus* pupating, and b) the number of days taken for *M. signatus* to pupate, in trial 2b. There were no significant differences between treatments in the number of lacewings pupating (contingency test, χ^2^=6.171, df=3, p=0.104, N=40), or the number of days taken for *M. signatus* to pupate (ANOVA, F_3,31_=1.06, p=0.379).
